# Greater strength of selection and higher proportion of beneficial amino acid changing mutations in humans compared to mice and *Drosophila melanogaster*

**DOI:** 10.1101/427583

**Authors:** Ying Zhen, Christian D. Huber, Robert W. Davies, Kirk E. Lohmueller

## Abstract

Quantifying and comparing the amount of adaptive evolution among different species is key to understanding evolutionary processes. Previous studies have shown differences in adaptive evolution across species; however, their specific causes remain elusive. Here, we use improved modeling of weakly deleterious mutations and the demographic history of the outgroup species and ancestral population and estimate that at least 20% of nonsynonymous substitutions between humans and an outgroup species were fixed by positive selection. This estimate is much higher than previous estimates, which did not correct for the sizes of the outgroup species and ancestral population. Next, we directly estimate the proportion and selection coefficients (*p*^*+*^ and *s*^*+*^, respectively) of newly arising beneficial nonsynonymous mutations in humans, mice, and *Drosophila melanogaster* by examining patterns of polymorphism and divergence. We develop a novel composite likelihood framework to test whether these parameters differ across species. Overall, we reject a model with the same *p*^*+*^ and *s*^*+*^ of beneficial mutations across species, and estimate that humans have a higher *p*^*+*^*s*^*+*^ compared to *D. melanogaster* and mice. We demonstrate that this result cannot be caused by biased gene conversion or hypermutable CpG sites. In summary, we find the proportion of beneficial mutations to be higher in humans than in *D. melanogaster* or mice, suggesting that organismal complexity, which increases the number of steps required in adaptive walks, may be a key predictor of the amount of adaptive evolution within a species.

## INTRODUCTION

Since the inception of molecular population genetics, there has been tremendous interest in quantifying the amount of adaptive evolution in different organisms. The neutral theory of molecular evolution postulated that beneficial mutations are rare, and that many of the substitutions between species are neutral (Kimura 1983). One early challenge to this theory originated from a comparison of polymorphisms and divergence at synonymous and nonsynonymous sites in *Drosophila* (Fay et al. 2002; Smith and Eyre-Walker 2002). Under models without positive selection, the ratio of nonsynonymous to synonymous changes should remain equal when comparing polymorphisms (*i.e.* differences within species) and divergence (*i.e.* differences between species). In contrast to this prediction, a genome-wide excess of nonsynonymous divergence between species was observed, a pattern indicative of an abundance of positive selection in *Drosophila*. More formally, Smith and Eyre-Walker (2002) proposed a statistic, α, which is the proportion of nonsynonymous substitutions between species that can be attributed to positive selection. Application of their approach in *Drosophila* has found that at least 40% of nonsynonymous substitutions have been fixed by positive selection (Smith and Eyre-Walker 2002).

Since the publication of the original study, α has been estimated from different species across the tree of life (Fay 2011). Estimates of α vary tremendously across species, tending to be higher in insects (Andolfatto 2005; Eyre-Walker and Keightley 2009), but much lower in primates and plants (Boyko et al. 2008; Eyre-Walker and Keightley 2009; Gossmann et al. 2010). In these latter species, formal tests have been unable to reject the hypothesis that α is zero (*i.e.* no positive selection; Boyko et al. 2008; Foxe et al. 2008; Eyre-Walker and Keightley 2009). It is not clear why α varies across species. One possibility is that α is higher for species with larger population sizes, which could occur if adaptation is mutation limited. Here, species with larger population sizes would have a higher rate of beneficial mutations. The fixation probability of a given beneficial mutation also would be higher in species with larger population size, but this effect is likely to only be important for very weakly beneficial mutations. Evidence indicates that, in some cases, α is indeed correlated with population size. For example, Phifer-Rixey et al. (2012) found that estimates of α were higher for species of mice that have larger population sizes compared to species with smaller population sizes. Further, studies found a positive correlation between α and population size when comparing different species of sunflowers (Strasburg et al. 2011) and from phylogenetically diverse taxa (Gossmann et al. 2012). More recently, Galtier (2016) found a positive correlation between α and effective population size for 44 animal species. Additional evidence that there is more positive selection in larger populations stems from recent analyses of linked selection. Corbett-Detig et al. (2015) found increased evidence for linked selection in species with larger population sizes, though the mechanism driving this pattern is not immediately clear. Further, Nam et al. (2017) have suggested that across primates, species with larger population sizes have had more selective sweeps.

While this evidence suggests that adaptation could be mutation limited and that this could be driving the variation in α across species, it is important to note that other factors could be influencing the α statistic (Rousselle et al. 2018; Messer and Petrov 2013). As the denominator of α is the total number of observed differences between species, it is sensitive to the fixation of weakly deleterious mutations. For two populations with the same number of beneficial substitutions, the one with a higher number of substitutions due to weakly deleterious mutations will have a lower α. Indeed, because the number of fixed weakly deleterious mutations is inversely related to population size (Ohta 1973, 1992), this effect could drive the correlation between α and population size. In support of this prediction, Galtier (2016) found that the rate of adaptive divergence (omega-a), which does not depend on the total number of divergent sites between species (Gossmann et al. 2012), showed no correlation with population size. Similar arguments have been made by Phifer-Rixey et al. (2012). Further, for humans, it has been suggested that α had been underestimated due to the presence of weakly beneficial mutations still segregating as polymorphisms (Galtier 2016; Uricchio et al. 2019). Indeed, methods that account for weakly beneficial mutations still segregating as polymorphisms infer slightly higher values of α = 0.24 between humans and chimpanzee, and α = 0.135 in the human lineage (Galtier 2016; Uricchio et al. 2019). In addition, population sizes of the outgroup and ancestral population determine the rate of fixation of weakly deleterious mutations in the outgroup lineage and in the ancestral population, and could potentially influence the estimate of α (Rousselle et al. 2018; McDonald and Kreitman 1991; Eyre-Walker 2002).

Other studies quantified positive selection by focusing on the proportion of beneficial mutations (*p*^*+*^) and their selection coefficients (*s*^*+*^). Boyko *et al.* (2008) found that by assuming a fraction (0 to 1.86%) of new mutations is positively selected, they could better match the frequency spectrum of polymorphisms and the counts of human-chimpanzee divergence. Models with weaker selection coefficients for beneficial mutations tended to have a higher proportion of positively selected mutations than models with stronger selection (Boyko et al. 2008). Several studies also have estimated *p*^*+*^ and *s*^*+*^ in *Drosophila* species and mice (Sella et al. 2009; Schneider et al. 2011; Elyashiv et al. 2016; Campos et al. 2017; Keightley et al. 2016; Booker and Keightley 2018). However, there has not been a systematic comparison across species.

Here we compare the amount of adaptive evolution in primates, rodents, and *Drosophila*. We use two complementary approaches that quantify different aspects of the adaptive process. First, we use improved modeling of weakly deleterious mutations and demographic models, particularly correcting for the sizes of outgroup species and ancestral population to infer α. Second, we estimate *p*^*+*^ and *s*^*+*^ of newly arising beneficial mutations by examining patterns of polymorphism and divergence. We develop a composite likelihood framework to test whether these parameters differ across taxa. This approach enables a more direct comparison of the amount of beneficial mutations across species and is less confounded by the fixation of weakly deleterious mutations.

## RESULTS

### Estimates of α for multiple species using the MK method

We first estimated α from coding regions of primates, rodents, and *Drosophila*. We analyzed published genomic datasets to obtain counts of synonymous and nonsynonymous polymorphisms (*P*_*S*_ and *P*_*N*_) and divergent sties between species (*D*_*S*_ and *D*_*N*_). For computation of α in humans, we used chimpanzee and macaque as outgroup species. For mice and *D. melanogaster*, we used rat and *D. simulans* as the outgroup species, respectively. In total, 19.1 Mb of coding sequence for primates, 26.6 Mb of coding sequence for rodents, and 15.8 Mb of coding sequence for *Drosophila* were used in our analysis (see Methods).

An extension of the McDonald-Kreitman test was used to estimate α (Smith and Eyre-Walker 2002; Table 1; Supplemental Table S1). To examine the effect of slightly deleterious mutations on α, we filtered the data with several minor allele frequency (MAF) cutoffs (Messer and Petrov 2013; Fay et al. 2001; Charlesworth and Eyre-Walker 2008) (Supplemental Table S2). For example, after removing low frequency polymorphisms with MAF less than 20%, the estimated α is close to zero for the human-chimpanzee comparison (Table 1), consistent with previous estimates (Boyko et al. 2008; Eyre-Walker and Keightley 2009). However, for the human-macaque comparison, α is -0.22 (Table 1), suggesting that the choice of outgroup species could greatly influence α. Nevertheless, these results suggest at most only a very small proportion of nonsynonymous substitutions have been fixed by positive selection in primates. In contrast, for the *D. melanogaster-D. simulans* and mouse-rat comparisons, the estimated α is 49% and 40%, respectively, with MAF filter at 20% (Table 1). Both of these estimates are comparable to previous studies (Andolfatto 2005; Eyre-Walker and Keightley 2009; Phifer-Rixey et al. 2012). These results suggest that the proportion of substitutions fixed by positive selection varies drastically across species, and for taxa with larger population sizes, like rodents and *Drosophila*, adaptive forces may have had a greater contribution to divergence.

**Table 1.** Estimates of α using different methods

|  |  |  | Method of Inference |  |  |  |  |  |
| --- | --- | --- | --- | --- | --- | --- | --- | --- |
| | Species | Outgroup | MK-method:<br>all | MK-method:<br>MAF>20% | Model-based:<br>Simple Model | Model-based:<br>Complex Model | DFEα:<br>$\alpha$ | DFEα:<br>$\omega_a$ |
| Full data | Human | Chimpanzee | -0.41 | -0.01 | 0.11 | 0.25 | 0.24 | 0.07 |
|  | Human | Macaque | -0.70 | -0.22 | -0.08 | 0.26 | 0.02 | 0.01 |
|  | Human lineage | - | - | - | 0.06 | 0.16 | - | - |
|  | *D. melanogaster* | *D. simulans* | -0.13 | 0.49 | 0.53 | 0.60 | 0.71 | 0.14 |
|  | Mice | Rat | 0.25 | 0.40 | 0.45 | 0.41 | 0.51 | 0.12 |
| SSWW only | Human | Chimpanzee | -0.37 | 0.08 | 0.13 | 0.26 | - | - |
|  | Mice | Rat | -0.14 | 0.08 | 0.20 | 0.10 | - | - |

### Model-based inference of α

Model-based approaches estimate α by contrasting the observed number of nonsynonymous differences between species *D*_*NO*_ with the number expected under a demographic model and a distribution of fitness effects (DFE) including only neutral and deleterious mutations *D*_*NE*_ (Boyko et al. 2008; Eyre-Walker and Keightley 2009). The excess of observed *D*_*NO*_ compared to the predicted *D*_*NE*_ is attributed to fixations driven by positive selection. These methods also assume that the population sizes of outgroup species and the ancestral population of the ingroup and outgroup are the same as the ancestral size of the ingroup population (*Nanc.in*). We refer to this demographic model as the Simple model (Fig. 1A).

**Figure 1.**
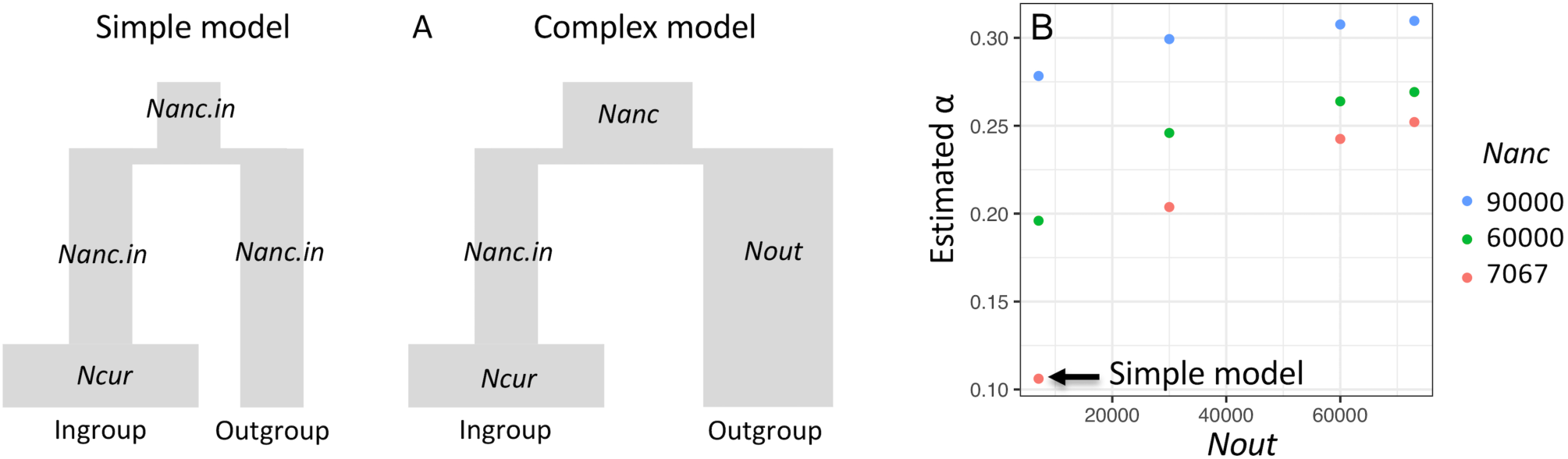
The ancestral and outgroup population sizes greatly influence α. (A). Schematic demographic models illustrate the Simple and the Complex model with associated parameters. In the Simple model, the size of the ancestral population (*Nanc)* as well as the size of the outgroup (*Nout*) are assumed to be the same as the ancestral size of the ingroup (*Nanc.in*). This assumption is relaxed in the Complex model. (B). Effect of *Nanc* and *Nout* on estimates of α for humans using the chimpanzee as an outgroup. Colors denote different values of *Nanc*. Arrow points to the estimate of α from the Simple model where *Nanc = Nout = Nanc.in* = 7067.

Using the Simple model, we first estimated the divergence time that best fit our data using the observed *D*_*S*_. We then predicted *D*_*NE*_ using this estimated divergence time, the gamma-distributed DFE inferred in Huber et al. (2017), and the demographic model of the ingroup species (Supplemental Text; Supplemental Table S3 and S4). We then estimated that α=11% for humans and chimpanzee. This estimate is comparable to the inference by Boyko *et al.*, where authors implemented a similar approach and estimated that approximately 10% of human-chimpanzee nonsynonymous substitutions were fixed by positive selection (Boyko et al. 2008). When using macaque as the outgroup species, we estimate that α is negative (Table 1). While it is possible that this difference could reflect distinct evolutionary events experienced by different outgroups, it could also be an artifact of the modeling assumptions. Previous estimates of the human-chimpanzee ancestral population size (*Nanc*) range from 12,000 to 125,000 using different datasets and methods (Chen and Li 2001; Hobolth et al. 2007; Burgess and Yang 2008; Prado-Martinez et al. 2013; Schrago 2014). The estimated chimpanzee population size (*Nout)* is around 30,000 (Prado-Martinez et al. 2013; Hvilsom et al. 2012; Fischer et al. 2004) (Supplemental Table S5). However, the ancestral size inferred for the human lineage using human polymorphism data is 7067, which is much smaller than all previous estimates of *Nanc* and *Nout*. Using values of *Nanc* and *Nout* that are too small likely biases estimates of α because more of the nonsynonymous substitutions are incorrectly attributed to the fixation of weakly deleterious mutations, causing α to be under-estimated.

To more accurately model the larger *Nanc* and *Nout*, we use the Complex model which allows *Nout* and *Nanc* to differ from *Nanc.in* (Fig. 1A). For humans, the larger *Nanc* is very ancient, thus it does not affect the polymorphism pattern within humans (confirmed by coalescent simulations, see Supplemental Text). We calculated the number of substitutions fixed in the ingroup, outgroup, and ancestral populations. The total expected number of differences between a pair of species is the sum of differences at all three stages. We first explored how *Nout* and *Nanc* affect the inferred α. For the human-chimpanzee comparison, the inferred α changes dramatically with different values of *Nout* and *Nanc*. Larger values result in larger estimates of α (Fig. 1B). For example, when *Nout* = 30,000 (Prado-Martinez et al. 2013; Hvilsom et al. 2012; Fischer et al. 2004) and *Nanc* = 60,000 (Chen and Li 2001; Hobolth et al. 2007; Prado-Martinez et al. 2013), supported by many previous studies (Supplemental Table S5), α is approximately 24.6%. When using other values of *Nout* and *Nanc* within the ranges of previous estimates, the corresponding estimates of α may differ, but remain above 20% (Fig. 1B). We next revisited the human-macaque comparison under the Complex model (Supplemental Fig. S1), using *Nout* = 73,000 (Hernandez et al. 2007; Xue et al. 2016), *Nanc =* 48,000 (McVicker et al. 2009), and changing the ingroup population size to 60,000 at human-chimpanzee divergence time to match what we modeled in human-chimpanzee (Supplemental Table S4). Here we infer approximately 26.0% of human-macaque nonsynonymous substitutions were fixed by positive selection which is comparable to what we found for the human-chimpanzee analysis. These estimates of α for primates using the more realistic Complex demographic model are much higher than previous estimates, implying that there is a greater contribution of positive selection to nonsynonymous divergence than previously appreciated.

Similarly, we estimated α for *Drosophila* and rodents using the Complex demographic model. For *Drosophila*, it had been inferred that *D. simulans* have slightly larger *N*_*e*_ than *D. melanogaster* (Andolfatto et al. 2011), so we set *Nout* to be 1.5× the current population size of *D. melanogaster* (*Ncur*) at the species’ split (Supplemental Table S4). For mice, a previous study estimated that the outgroup rat species has an effective population size about fivefold lower than wild house mice (Ness et al. 2012). Thus, we set the *Nout* to be 0.2× *Ncur* of mice (Supplemental Table S4). Since there is limited knowledge of the population sizes of the ancestor of *D. melanogaster* and *D. simulans*, and ancestor of mice and rat, for these two comparisons, we assume *Nanc = Nout.* Using these Complex models for *D. melanogaster-D.simulans* and mouse-rat, we estimated their α to be 60% and 41%, compared with 53% and 45% using the Simple model, respectively (Table 1). The differences between these estimates reflect the importance of accurately modeling the population size of outgroup species for calculations of α.

We estimated α for substitutions that occurred exclusively on the human lineage, using the human-macaque alignment to polarize sequence differences between human and chimpanzee. We estimate that α = 6.2% using the Simple model, and 16.0% using the Complex model (Table 1).

To compare our estimates of α to those from another model-based method that assumes *Nanc=Nout*=*Nanc.in*, we used *DFE-alpha*, to infer α for our three species pairs. Using this method, α is estimated to be 24% and 2% for humans using chimpanzee and macaque as outgroup respectively, 71% for *D. melanogaster-D. simulans*, and 51% for mouse-rat (Table 1). These estimates are all higher compared to estimates from Simple models. However, the estimates of α for primates differ significantly depending on whether the macaque or chimpanzee is used as the outgroup for humans.

### Testing whether p^+^ and s^+^ differ across species

We next estimated the DFE including new beneficial mutations. Our model of the DFE includes two additional parameters compared to the base model that only includes deleterious mutations. For each species, we estimate the proportion of new mutations that are beneficial (*p*^*+*^) and their selection coefficient (*s*^*+*^). We then test whether these two parameters differ across species.

The number of nonsynonymous differences between a pair of species (*D*_*N*_) is assumed to be Poisson-distributed (Sawyer and Hartl 1992), with rate parameter equal to:

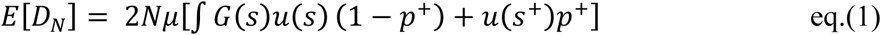

where *G*(*s*) is the DFE of deleterious and neutral mutations from Huber et al. (2017), *u*(*s*) is the fixation probability (Kimura 1962) of deleterious and neutral mutations, *u*(*s*^+^) is the fixation probability of beneficial mutations, and *p*^*+*^ is the proportion of new mutations that are beneficial. We then use a Poisson log-likelihood function for *D*_*N*_ in each species and a series of likelihood ratio tests (LRTs) to determine whether *p*^*+*^ and *s*^*+*^ differ across primates, rodents, and *Drosophila* (see Methods).

Using this framework, we find that the full model H1, where each taxon is allowed to have its own *p*^*+*^ and *s*^*+*^, fits *D*_*N*_ significantly better than the constrained null model, where *p*^*+*^ and *s*^*+*^ are constrained to be the same across all three taxa (LRT statistic Λ=124,974, df=4, *P*<10^−16^; Fig. 2; Supplemental Fig. S2; Supplemental Table S6).

**Figure 2.**
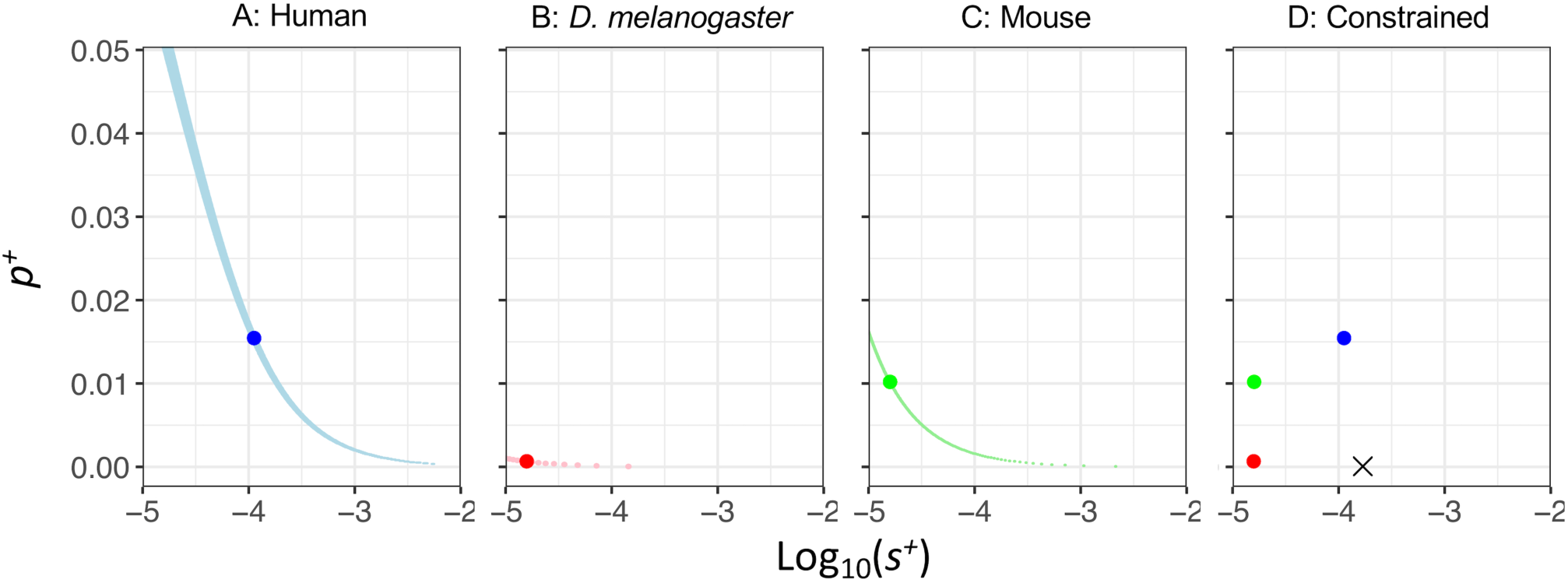
Log-likelihood surfaces for *p*^*+*^ and *s*^*+*^ for different species. (A) Human. (B) *D. melanogaster*. (C) Mouse. (D) The constrained model (H0), where *p*^*+*^ and *s*^*+*^ are constrained to be the same across all three taxa. Log-Likelihoods (LL) are calculated using grid search method of log_10_(*s*^*+*^) in the range of -5 to -2 and *p*^*+*^ in the range of 0-7.5%. Blue denotes human, red denotes *D. melanogaster*, and green denotes mouse. The large points represent the MLE for each species, the black cross in panel D represents the MLE of the constrained model, and the lighter colors show grid points within 3 LL units of each MLE. The Complex model is used for each species and we use the chimpanzee as the outgroup for humans.

We estimate that humans have a high2er proportion of beneficial nonsynonymous mutations than *D. melanogaster* and mice (Supplemental Table S6 and Fig. 2). Specifically, we estimate that approximately 1.55% of new nonsynonymous mutations in humans are beneficial with *s*^*+*^ of approximately 1.12×10^−4^ (outgroup: chimpanzee), approximately 0.0675% of new nonsynonymous mutations in *D. melanogaster* are beneficial with *s*^*+*^ of approximately 1.58×10^−5^, and approximately 1.02% of new nonsynonymous mutations in mice are beneficial with *s*^*+*^ of 1.60×10^−5^ (Supplemental Table S6). Models with a larger *s*^*+*^ tend to have a lower proportion of positively selected mutations than models with smaller *s*^*+*^. Consequently, the likelihoods of these parameter values can be very close to the likelihoods at the MLEs. We also compared *p*^*+*^ and *s*^*+*^ between pairs of taxa (*i.e.* primate *vs.* rodent; primate *vs. Drosophila*; rodent *vs. Drosophila*). In all pairwise tests (regardless of outgroup or demography), the model where each species has its own *p*^*+*^ and *s*^*+*^ fits the observed *D*_*N*_ significantly better than a model where *p*^*+*^ and *s*^*+*^ are constrained to be the same in the tested two taxa (Supplemental Table S6).

We next investigated whether it is possible that either *p*^*+*^ or *s*^*+*^ is the same across taxa but the other parameter varies. Specifically, we allowed *p*^*+*^ to differ across primates, rodents, and *Drosophila*, then explored whether a model with the same *s*^*+*^ could fit all species. This is shown in conditional likelihood plots, where assuming the same *s*^*+*^ for all species, humans would need a higher proportion of beneficial mutations compared to mice and *D. melanogaster* to match the observed *D*_*N*_ (Fig. 3A). Similarly, allowing *s*^*+*^ to differ across species, a model with the same *p*^*+*^ across all species could fit the data. When we forced the same *p*^*+*^ for all species, *s*^*+*^ for beneficial mutations in humans would be larger compared to that in mice and *D. melanogaster* (Fig. 3B). However, for the same *p*^*+*^ value, *s*^*+*^ could not be the same in all species. Similarly, for the same *s*^*+*^ value, *p*^*+*^ could not be the same across species.

**Figure 3.**
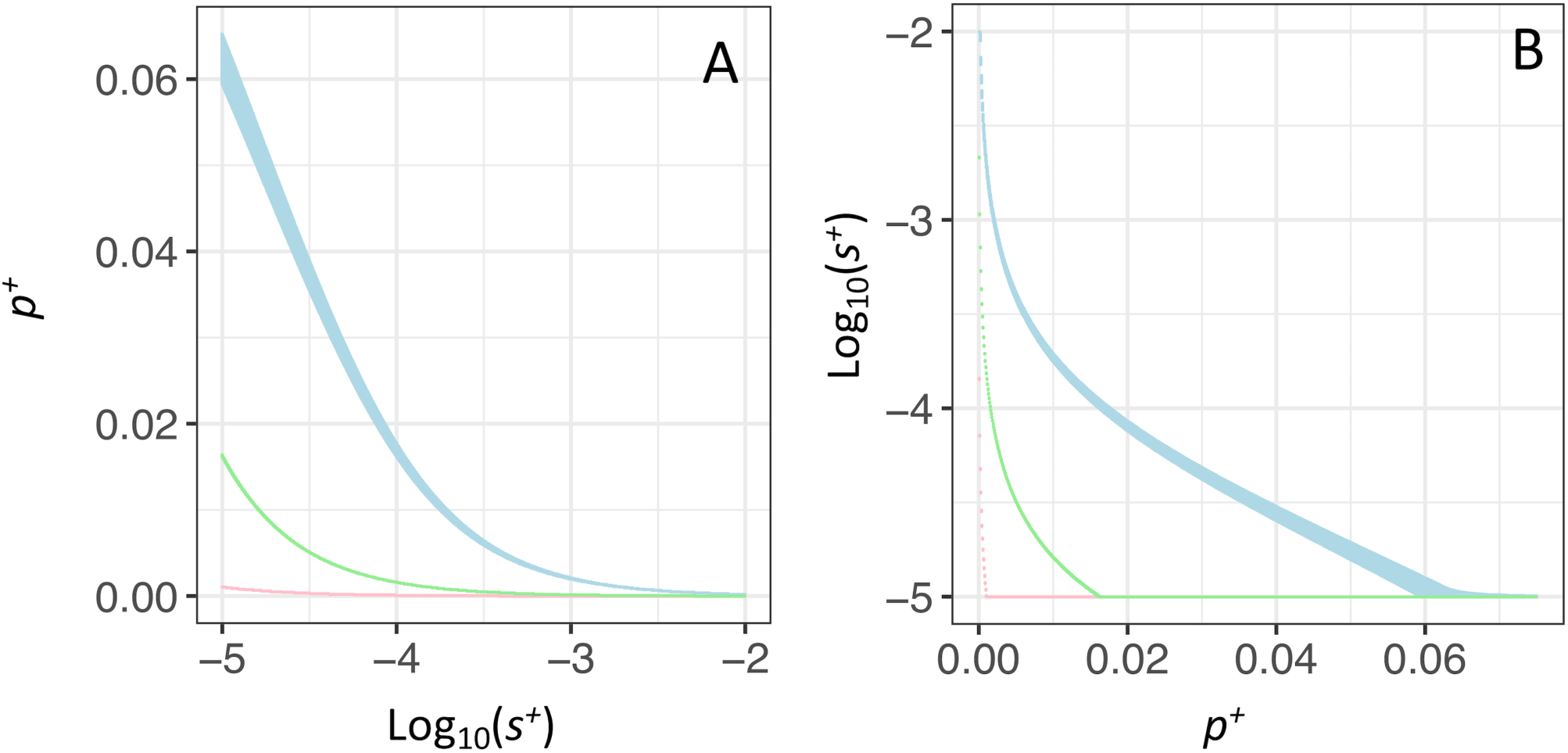
Conditional log-likelihood surfaces. (A) Maximizing *p*^*+*^ given particular values of *s*^*+*^ and (B) maximizing *s*^*+*^ given particular values of *p*^*+*^. Only grid points within 3 LL units of the MLEs for each parameter for each species are shown. Light blue denotes human, pink denotes *D. melanogaster*, and light green denotes mouse.

Motivated by the observation that larger values of *p*^*+*^ and smaller values of *s*^*+*^ can fit the data as well as models with smaller values of *p*^*+*^ and larger values of *s*^*+*^, we also estimated a composite parameter *p*^*+*^*s*^*+*^, the product of *p*^*+*^ and *s*^*+*^. We find clear evidence that regardless of the model of ancestral demography or outgroup, humans have a significantly higher *p*^*+*^*s*^*+*^ than do *D. melanogaster* and mice (Fig. 4). This can be seen by looking at the log-likelihood curves, which are quite peaked, with little overlap across species. Thus, approximate 95% confidence intervals on *p*^*+*^*s*^*+*^ for the different species do not overlap, implying more positive selection in humans than in *D. melanogaster* and mice.

**Figure 4.**
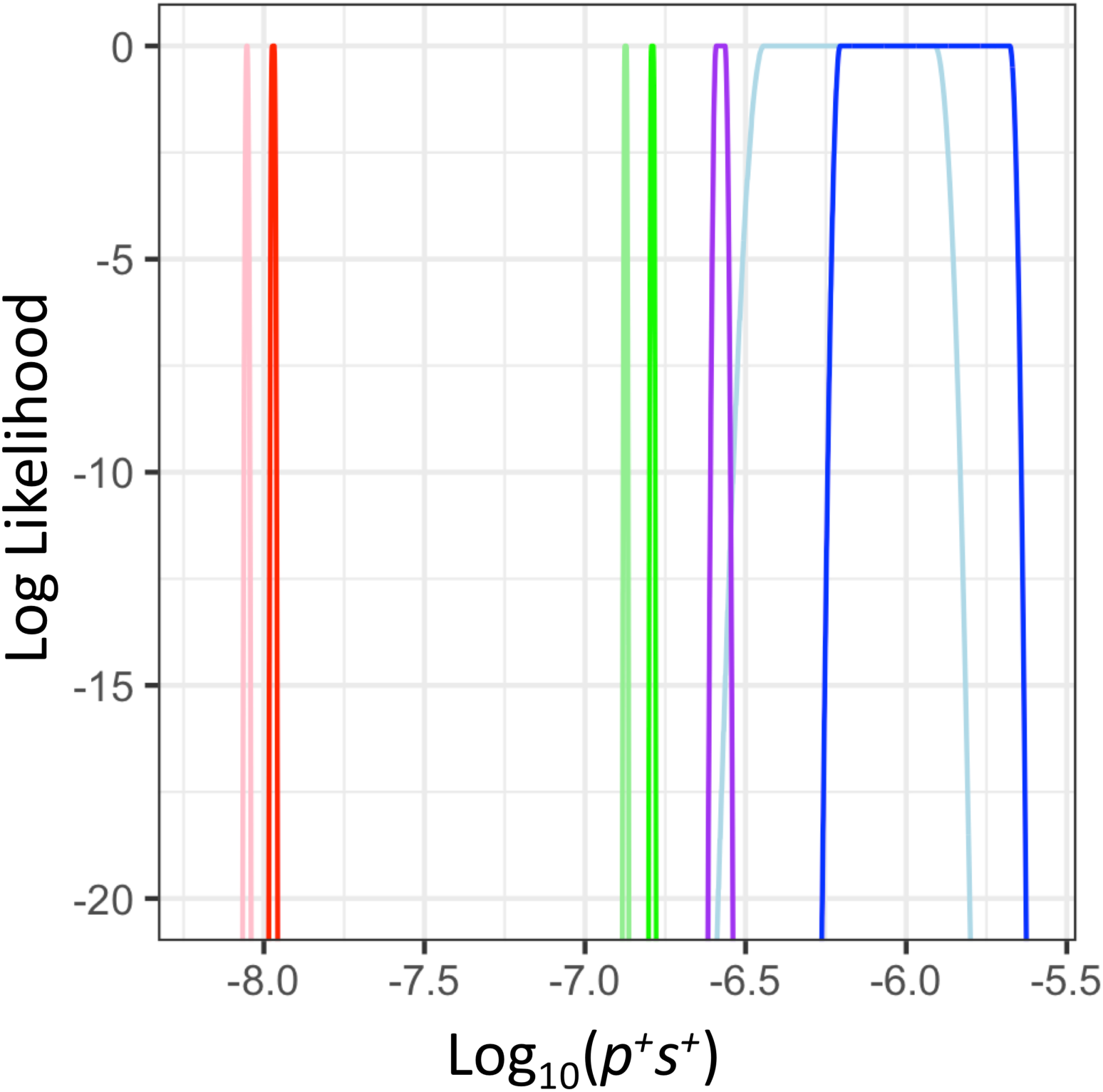
The composite parameter *p*^*+*^*s*^*+*^, capturing the proportion of beneficial mutations and the strength of selection, differs across species. Log-likelihood curves for *p*^*+*^*s*^*+*^ in the three species. Red denotes the inference for *D. melanogaster*, green denotes the inference for mouse, blue denotes the inference for human using the chimpanzee as the outgroup, and purple denotes the inference for human using the macaque as the outgroup. Lighter colors denote the Simple model. Darker colors denote the Complex model that better models the ancestral demography and population size of the outgroup. Note that regardless of which demographic model is used, the log-likelihood curves from the different species do not overlap within the top 500 log-likelihood units, suggesting *p*^*+*^*s*^*+*^ is significantly different among taxa.

### Testing whether γ+ and p+ differ across species

Humans, *D. melanogaster*, and mice have drastically different population sizes, which can influence the efficacy of selection within each species. Thus, we next examined whether the selection coefficient scaled by current population size (***γ***^*+*^=*2Ns*^*+*^) and *p*^*+*^ differ across primates, rodents, and *Drosophila*.

We find the model (Full model H1) where each taxon has its own different ***γ***^*+*^ and *p*^*+*^ fits the observed *D*_*N*_ significantly better than a model (constrained model H0) where ***γ***^*+*^ and *p*^*+*^ are constrained to be the same across all three taxa (LRT statistic Λ=3,541; df=4, *P*<10^−16^, Fig. 5; Supplemental Fig. S2; Supplemental Table S6).

**Figure 5.**
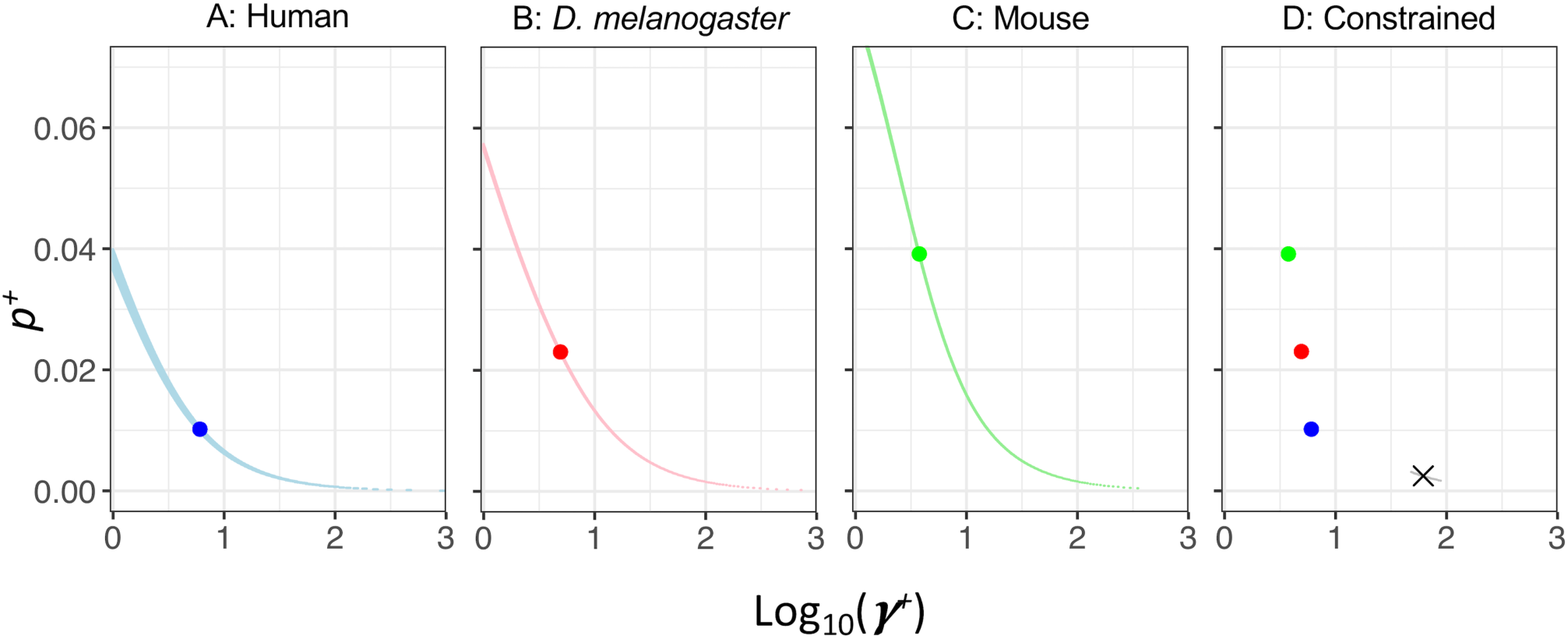
Log-likelihood surfaces for *p*^*+*^ and *γ*^*+*^ for different species. (A) Human. (B) *D. melanogaster*. (C) Mouse. (D) The constrained model, H0, where *p*^*+*^ and ***γ***^*+*^ are constrained to be the same across all three taxa. Log-likelihoods are calculated using grid search method of log_10_(***γ***^*+*^) in the range of 0-3 and *p*^*+*^ in the range of 0-7.5%. Blue denotes human, red denotes *D. melanogaster*, and green denotes mouse. The large points represent the MLE for each species, the black cross in panel D represents the MLE of the constrained model, and the lighter colors show grid points within 3 LL units of each MLE. The Complex model is used for each species and we use chimpanzee as the outgroup for humans.

Under the full model, we estimate that approximately 1% of new nonsynonymous mutations in humans are beneficial with ***γ***^*+*^ of 6.05, approximately 2% of new nonsynonymous mutations in *D. melanogaster* are beneficial with ***γ***^*+*^ of 4.92, and approximately 4% of new nonsynonymous mutations in mice are beneficial with ***γ***^*+*^ of 3.76 (Supplemental Table S6). Models with larger ***γ***^*+*^ tended to have a lower proportion of positively selected mutations than models with smaller ***γ***^*+*^, and the likelihoods can be very close to those at the MLEs. The current population sizes are 16,539, 7,616,700, 488,948 for humans, *D. melanogaster* and mice, respectively. As a result, we are searching for *p*^*+*^ within the same range of ***γ***^*+*^ (see Methods), but markedly different ranges of *s*^*+*^ for these three species. Thus, the relative ordering of the MLEs of *p*^*+*^ across species considering ***γ***^*+*^ is not necessarily the same as that for the *s*^*+*^ results described above.

We also compared ***γ***^*+*^ and *p*^*+*^ between pairs of taxa. We find that for the human-*D. melanogaster* and human-mice pairs, the models where each species has its own different ***γ***^*+*^ and *p*^*+*^ fit the observed *D*_*N*_ significantly better than a model where ***γ***^*+*^ and *p*^*+*^ are constrained to be the same in the tested two taxa (Supplemental Table S6). This result is robust regardless of outgroup species. However, we cannot reject the hypothesis that mice and *D. melanogaster* have same ***γ***^*+*^ and *p*^*+*^ (Supplemental Table S6).

We next investigated whether it is possible that either ***γ***^*+*^ or *p*^*+*^ is the same across taxa, while the other parameter varies. With the same ***γ***^*+*^ for all species, humans would need a lower proportion of beneficial mutations compared to mice and *D. melanogaster* to fit the observed *D*_*N*_ (Supplemental Fig. S3A). When we constrain *p*^*+*^ to be the same for all taxa, humans have a smaller ***γ***^*+*^ for beneficial mutations (Supplemental Fig. S3B). However, for the same *p*^*+*^ value, ***γ***^*+*^ cannot be the same in all taxa. Similarly, for the same ***γ***^*+*^ value, *p*^*+*^ cannot be the same across taxa. We also estimated a composite parameter *p*^*+*^***γ*** ^*+*^, the product of *p*^*+*^ and ***γ*** ^*+*^. We find that *p*^*+*^***γ***^*+*^ is not as distinct across different taxa as is *p*^*+*^*s*^*+*^. In the Complex models, humans have a smaller *p*^*+*^***γ*** ^*+*^ than mice. However, the log-likelihood curves of *p*^*+*^***γ*** ^*+*^ overlap between *D. melanogaster* and mice, and between *D. melanogaster* and humans, suggesting they are not significantly different (Supplemental Fig. S4).

### *Effects of biased gene conversion and hypermutable* CpG *sites*

Biased gene conversion (BGC) is the preferred transmission of G/C alleles (S: strong alleles) at the expense of A/T alleles (W: weak alleles). This process is common in mammals and could impact patterns of genetic diversity (Duret and Galtier 2009; Lachance and Tishkoff 2014; Bolívar et al. 2015) and bias estimates of rate of positive selection (Corcoran et al. 2017), especially when comparing between taxa with BGC and taxa without BGC, *i.e. D. melanogaster* (Robinson et al. 2014). Further, methylated CpG sites in mammals have a higher mutation rate to T alleles due to deamination of the C nucleotide (Sved and Bird 1990; Duncan and Miller 1980), which could possibly bias inferences of selection.

To test whether BGC and hypermutable CpG sites drive the observed pattern of positive selection across primates, rodents, and *Drosophila*, we filtered primate and rodent data to keep only strong to strong or weak to weak mutations (herein called SSWW mutations), which are not affected by BGC and are not CpG changes (Supplemental Text, Supplemental Fig. S5). We then re-inferred α. For primates, α is comparable to that without filtering (Table 1). For rodents, however, α is substantially lower than that before filtering, suggesting that biased gene conversion and CpG mutational processes may account for some of the nonsynonymous differences between mouse and rat.

Removing the effect of BGC and CpG sites in primates and rodents, we again quantify and compare the strength and proportion of new beneficial mutations across all three taxa. The model where each taxon has its own *p*^*+*^ and *s*^*+*^ fits the observed *D*_*N*_ significantly better than a model where *p*^*+*^ and *s*^*+*^ are constrained to be the same across all three taxa (LRT statistic Λ=825, df=4, *P*<10^−16^; Supplemental Fig. S6; Supplemental Table S6). Models comparing each pair of taxa (*e.g.* primates *vs.* rodents; primates *vs. Drosophila*; rodents *vs. Drosophila*) suggest that each taxon has its own unique *p*^*+*^ and *s*^*+*^, regardless of demography (Supplemental Table S6). Allowing *p*^*+*^ to differ across taxa, a model with the same *s*^*+*^ across all taxa could fit the data, and *vice versa.* When we force the same *s*^*+*^ for all species, humans still have the highest proportion of new beneficial mutations (Supplemental Fig. S6E). Constraining *p*^*+*^ to be the same for all species, humans have the largest selection coefficient (Supplemental Fig. S6F).

Similarly, the model where each taxon has its own *p*^*+*^ and ***γ***^*+*^ fits the observed *D*_*N*_ significantly better than a model where *p*^*+*^ and ***γ***^*+*^ are constrained to be the same across all three taxa (LRT Λ=4496, df=4, *P*<10^−16^; Supplemental Table S6 and Fig. S7). Models comparing each pair of taxa suggest that each taxon has its own unique *p*^*+*^ and ***γ***^*+*^, regardless of demography (Supplemental Table S6). Allowing *p*^*+*^ to differ across species, a model with the same ***γ***^*+*^ across all taxa could fit the data, and *vice versa*. When we constrain ***γ***^*+*^ to be the same for all species, *D. melanogaster* have the highest proportion of new mutations being beneficial, and mice have the lowest proportion of new mutations that are beneficial. When we constrain *p*^*+*^ to be the same for all species, *D. melanogaster* have the largest ***γ***^*+*^, and mice have the smallest ***γ***^*+*^(Supplemental Fig. S7).

Lastly, we estimated two composite parameters, *p*^*+*^*s*^*+*^ and *p*^*+*^***γ*** ^*+*^ for SSWW mutations. Humans again have a significantly higher *p*^*+*^*s*^*+*^ than *D. melanogaster* and mice (Supplemental Fig. S8), suggesting more positive selection in humans than in *D. melanogaster* and mice. Similar to the full dataset, *p*^*+*^***γ*** ^*+*^ is not as distinct across different species as *p*^*+*^*s*^*+*^. Under the Complex model, we find that humans and *D. melanogaster* have a significantly (*i.e.* the log-likelihood curves do not overlap within 2 units of the MLEs) larger *p*^*+*^***γ*** ^*+*^ than mice, and the log-likelihood curves of *p*^*+*^***γ*** ^*+*^ for humans and *D. melanogaster* overlap near the MLEs (Supplemental Fig. S8).

## DISCUSSION

We have quantified the amount of adaptive evolution in multiple species with differing degrees of complexity and population size using two different approaches. First, we examined the proportion of nonsynonymous substitutions fixed by positive selection and found that this proportion is higher in *Drosophila* than in rodents and primates, consistent with prior work (Andolfatto 2005; Eyre-Walker and Keightley 2009; Boyko et al. 2008; Fay 2011). However, after correcting for the population sizes of the outgroup and ancestral population, we infer that α is much higher in primates than what was inferred in previous studies. Second, we show that the DFE of new beneficial mutations differs across species. The species with the smaller population size and greater complexity (*i.e.* humans) has stronger and/or more abundant new beneficial mutations than the other two species with much larger population sizes (*i.e.* mice and *D. melanogaster*). These results are robust to the choice of outgroups, BGC, and hypermutable CpG sites.

One major improvement of our method to infer α over previous similar approaches from Boyko et al. and *DFE-alpha* (Boyko et al. 2008; Eyre-Walker and Keightley 2009) is that we allow the outgroup and ancestral population sizes to differ from the ancestral population size of the ingroup. In Boyko *et al.* (2008), the outgroup population size is assumed to be the same as that found in the ancestral ingroup population. *DFE-alpha* allows for inference of demographic parameters of one- to three-epoch models, and the outgroup population size is assumed to be the same as the ancestral ingroup population. When considering species like humans in relation to other primates, like chimpanzee or macaque, this assumption certainly does not hold as primate species have population sizes at least several fold larger than the estimated ancestral human population size of approximately 7,000 individuals (Chen and Li 2001; Hobolth et al. 2007; Burgess and Yang 2008; Prado-Martinez et al. 2013; Schrago 2014; Hernandez et al. 2007; McVicker et al. 2009; Hvilsom et al. 2012; Wall 2013; Xue et al. 2016). The sizes of the outgroup and ancestral population matter because they affect the fixation probability of weakly deleterious alleles (Kimura 1983; Ohta 1973, 1992). As such, the number of nonsynonymous substitutions attributed to weakly deleterious mutations is highly affected by the population sizes of the outgroup and ancestral population. Consequently, estimates of α are affected as well. Our approach includes a more complex demographic model that allows the ancestral and outgroup populations to have their own sizes (*Nanc* and *Nout*). By using these more realistic population sizes, the α estimates we obtained for primates are similar when using chimpanzee or macaque as outgroup species. This is strong evidence that our method is more accurate as all the other methods give drastically different estimates of α using these two different outgroups. Interestingly, Eyre-Walker and Keightley (2009) also suggested that α in humans could be as high as 0.31 if the effective population size of humans and macaques was much higher than 10,000 until very recently, foreshadowing our current estimates. Future studies of adaptive substitutions should carefully consider ancestral and outgroup population sizes and should use statistical methods that model them realistically.

Here we have quantified adaptive evolution from two different perspectives. First, we estimate the proportion of adaptive nonsynonymous substitutions between species, α. This statistic measures the endpoint where a number of factors such as demography, genetic drift, and natural selection all come into play. Second, we estimate the DFE of newly arising beneficial mutations, *i.e. p*^*+*^ and *s*^*+*^. This approach aims to understand the properties of new beneficial mutations, the beginning point where beneficial mutations appear and enter the population. Thus, it is possible that these two approaches can yield qualitative opposite results. Indeed, we have shown that *D. melanogaster* have the largest values of α, but the smallest *p*^*+*^*s*^*+*^.

We previously found 15% of nonsynonymous mutations in humans are weakly beneficial (Huber et al. 2017). Importantly, because Huber *et al.* only analyzed polymorphism data, they did not consider strongly beneficial mutations that became substitutions between species. Our present study leverages genome sequences of several related species pairs to estimate the proportion of strongly beneficial (*s*^*+*^>10^−5^) mutations that are fixed between species. Weakly beneficial mutations are already accounted for in our model using the DFE from Huber *et al.* (2017) as many weakly beneficial mutations are likely segregating as polymorphisms. Intriguingly, we find that humans have a higher proportion of strongly beneficial mutations than *D. melanogaster*, mirroring the qualitative trend seen for weakly beneficial mutations in Huber *et al*. (2017). In the context of human evolution, it has been shown that weakly beneficial mutations that are still segregating could result in an underestimate of α (Uricchio et al. 2019; Galtier 2016). Thus, the estimate of α in humans could be even higher if adaptive polymorphisms were taken into account. Alternatively, our current approach of modeling the complex demography in the outgroup and ancestral population should correct for some of the bias in inferring α due to weakly beneficial mutations still segregating as polymorphisms. The reason for this is that weakly beneficial mutations still segregating as polymorphisms will be accounted for in the parametric DFE fit to the site frequency spectrum (SFS; as discussed above in the context of Huber et al. 2017). A parametric DFE including only deleterious and neutral mutations will treat this class of weakly beneficial mutations as nearly neutral (Tataru et al. 2017), inflating the expected divergence that can be accounted for by non-adaptive forces. Modeling the complex demography reduces the number of fixations from nearly neutral mutations that are treated in the model to be non-adaptive, including those from segregating weakly beneficial mutations.

Our estimate of α for human lineage is 16.0% using the Complex model, which is comparable to the estimate of human-lineage α by Uricchio *et al.* (2019), despite the use of different analytical approaches. α is expected to be lower on the human lineage as compared to the chimpanzee or macaque lineage due to the higher proportion of weakly deleterious amino acid substitutions on the human lineage to their smaller population size (Ohta 1973, 1992). For *D. melanogaster*, using the Complex model we estimate *p*^*+*^***γ***^*+*^ to be ∼0.1 which is within a factor of 2 of a previous estimate of Keightley et al. (2016). For mice, using the Complex model, we estimate *p*^*+*^***γ***^*+*^ to be ∼0.15 which is somewhat greater than the value of ∼0.05 inferred in Booker and Keightley (2018). The differences in estimates could be due to the use of different methods and models, such as our detailed modeling of the ancestral and outgroup population sizes.

We make several key modeling assumptions. First, we assume that the DFE is the same between the ingroup and outgroup for each pair of species (*e.g.* humans and chimpanzee have the same DFE). This assumption could be relaxed by using only polymorphism data to infer the beneficial DFE (Tataru et al. 2017), though not including divergence to an outgroup species considerably reduces the power to infer positive selection (Booker 2019). Second, our inference of *p*^*+*^ and *s*^*+*^, as well as related approaches (Schneider et al. 2011; Tataru et al. 2017; Booker and Keightley 2018), make the assumption of selection starting from a single mutation, since we use the fixation probability of a mutation introduced as a single copy. Selection on standing variation (*i.e.* one type of soft sweep) might bias the parameter estimates, though the meaning of *p*^*+*^ and *s*^*+*^ are not as clear under models with soft sweeps. α, however, should be accurately estimated even as long as the beneficial mutation reaches fixation. Furthermore, as our approach only focuses on strongly beneficial mutations that contribute to divergence, it is not sensitive to types of selection that do not reach fixation. These include another type of soft sweeps (Pritchard et al. 2010), where multiple independent beneficial mutations in the same gene are selected simultaneously, but no mutation becomes fixed, as well as polygenic adaptation where beneficial alleles only slightly increase in frequency, without reaching fixation. A last limitation comes from the model itself. It has been suggested that models where *s*^*+*^ does not change over time may not model the complexity of adaptive walks, where the first beneficial mutation may have a more strongly beneficial effect on fitness than subsequent beneficial mutations (Gillespie 2004; Lourenço et al. 2013). Thus, our estimates of *p*^*+*^ and *s*^*+*^ should be interpreted in this context as the average values over longer time periods and over different genetic backgrounds, rather than literal values that have stayed constant over time. In fact, our findings show that *p*^*+*^ and *s*^*+*^ have changed over deep evolutionary time, pointing to their dynamic nature.

One previous explanation for varying estimates of α across species was that adaptation is mutation limited and there are more beneficial mutations in organisms with larger population sizes. This view was not supported by conceptual arguments by Gillespie (2004) who suggested that the rate of environment change will matter more than the population size in determining the rate of adaptive evolution. The simulation study by Lourenço et al. (2013) that considered a changing DFE over time in the context of Fisher’s geometric model found that the population size only weakly related to α. Instead, the rate at which the environment changed was an important predictor of the amount of adaptive evolution, as environmental shifts moved the population from the fitness optimum, creating the opportunity for new beneficial mutations. Further, Rousselle et al. (2019) recently found a weak negative correlation between the amount of adaptive evolution and the amount of genetic diversity in modern populations, which supports the models of Lourenco *et al.*(2013) and Gillespie (2004). However, not all studies are in agreement on the role of the environment as Connallon and Clark (2015) found that environmental heterogeneity reduces the fraction of beneficial mutations by inflating the standardized mutation size in Fisher’s geometric model. Lourenço et al. (2013) also found that organismal complexity, here defined as the number of phenotypes under selection, was a key predictor of the amount of adaptive evolution within species. Through a “cost of complexity”, more complex organisms have a harder time adapting to new environmental conditions due to the additional constraints imposed by the increased number of traits under selection. As such, adaptive walks require more beneficial mutations (Orr 1998).

Our results presented here are in broad agreement with the conceptual model of Lourenço et al. (2013). Specifically, we do not find that species with larger population sizes (*i.e. D. melanogaster*) have more beneficial mutations. Instead, we find that *p*^*+*^ (as well as *p*^*+*^*s*^*+*^) is higher in humans than in *D. melanogaster* or mice. Second, while it is hard to precisely define organismal complexity, previous work has found more protein-protein interactions in humans than in *Drosophila* (Valentine et al. 1994; Stumpf et al. 2008), suggesting that humans may be more complex than *Drosophila*. If this is the case, then our findings of a higher *p*^*+*^ (and *p*^*+*^*s*^*+*^) in humans than *D. melanogaster* and mice supports the arguments from Lourenço et al. (2013) that adaptive walks after an environmental shift are less efficient and require more steps (*i.e.* beneficial mutations) in more complex organisms, leading to higher *p*^*+*^ in complex organisms. Additionally, differences in the degree of environmental change across species could also contribute to the disparate inferences of *p*^*+*^*s*^*+*^ across taxa. While it is hard to say which species has experienced more environmental shifts, it has been suggested that those species with longer generation times, like primates, may experience more environmental change per generation than species with shorter generation times, like *Drosophila* or rodents (Rousselle et al. 2019; Romiguier et al. 2014). Our finding of higher *p*^*+*^ in humans than *Drosophila* supports this intriguing prediction.

## METHODS

### Polymorphism and divergence data for humans, mice, and D. melanogaster

For humans, we used polymorphism data from 112 individuals from Yoruba in Ibadan, Nigeria (YRI) from the 1000 Genomes Project (1000 Genomes Project Consortium 2012). Published genome alignments of human and chimpanzee (hg19/pantro4), and human and *Macaca mulatta* (hg19/rheMac3) were downloaded from the UCSC genome browser (http://hgdownload.soe.ucsc.edu/goldenPath/hg19/). For *D. melanogaster*, we used the Drosophila Population Genomics Project phase 3 data, including 197 African *D. melanogaster* lines from Zambia, Africa (Lack et al. 2015). For divergence, *D. melanogaster* and *D. simulans* genic alignments (Dmel v5/Dsim v2) were extracted from the multi-species alignments from Hu et al. (2013). Only autosomal regions were used in our analysis. Human and *D. melanogaster* polymorphism data were filtered and down-sampled to 100 chromosomes as described in Huber et al. (2017).

For mice, raw data (fastq) was downloaded for 10 *M. m. castaneus* individuals that were collected in the northwest Indian state of Himachal Pradesh (Halligan et al. 2010, 2013). Reads were mapped against mouse genome mm9 using bwa (Li and Durbin 2009) and stampy (Lunter and Goodson 2011). Duplicate reads were marked using Picard, and further pre-processing was done following GATK Best Practice guidelines (McKenna et al. 2010). Variants were called using the GATK UnifiedGenotyper and filtered using the GATK VQSR using Affymetrix Mouse Diversity Genotyping Array sites (Yang et al. 2009). We further filtered the dataset to only retain sites with a sample size of at least 16 chromosomes and down-sampled all sites with larger sample size to a sample size of 16 chromosomes using the hypergeometric probability distribution. Published genome alignments of mice and rat (mm9/rn5) were downloaded from UCSC (http://hgdownload.soe.ucsc.edu/goldenPath/mm9/vsRn5/axtNet/). For each species, polymorphism and divergence data were intersected, and only coding regions shared by both datasets were used in our analysis.

In total, 19.1Mb of coding sequences for primates, 26.6 Mb of coding sequences for rodents and 15.8Mb of coding sequences for *Drosophila* were included. The nonsynonymous and synonymous total sequence lengths (*L*_*NS*_, *L*_*S*_) were estimated using multipliers of *L*_*NS*_ = 2.85 × *L*_*S*_ in *D. melanogaster*, and *L*_*NS*_ = 2.31 × *L*_*S*_ in mammals from Huber et al. (2017). In these filtered coding sequences, we annotated synonymous and nonsynonymous sites in both polymorphism and substitution data for each species. Human variants were annotated using the SeattleSeq Annotation pipeline (http://snp.gs.washington.edu/SeattleSeqAnnotation138/). Mice and Drosophila variants were annotated using SnpEeff v3.6 using the mice NCBIM37.66 annotation database and the *D. melanogaster* BDGP5.75 annotation database, respectively. Sites that are annotated as near-splice, or loss of function were removed. The ratio of nonsynonymous/synonymous differences between human and chimp sequences in our dataset is approximately 0.65, which is consistent with several previous reports from different datasets (Bustamante et al. 2005; Torgerson et al. 2009; Enard et al. 2014).

From the down-sampled polymorphism data, we calculated the synonymous and nonsynonymous SFS, and used the folded SFS for all further inferences to avoid misidentification of the ancestral state.

### Model-based estimates of α

To implement a model-based approach to estimate α, for each pair of species, we need a demographic model and a DFE for neutral and deleterious mutations to predict the *D*_*NE*_ that is accounted for by neutral and deleterious mutations. For primates and *Drosophila*, we use demographic and DFE parameters from Huber *et al.* (2017; Supplemental Table S3). For rodents, we conducted our own inference of these parameters by summarizing the polymorphism data in mice by the folded SFS (Supplemental Text).

To compute the expected divergence between species, we computed the divergence accumulated in ingroup population, outgroup population and ancestral population separately and summed them up. The divergence in the ingroup and outgroup population is a function of divergence time, effective population size, mutation rate, and selection coefficient and was calculated according to equation 13 in Sawyer and Hartl (1992). We computed the expected divergence under a gamma DFE model by Monte Carlo integration using one million gamma distributed selection coefficients. The contribution of the ancestral population to divergence between two species was computed numerically based on the diffusion equation, using *prfreq* (Boyko et al. 2008) and assuming the same gamma DFE. These calculations are implemented in our program *predicDiv* (https://github.com/LohmuellerLab/predicDiv). For both the Simple and the Complex models, we first estimate the divergence times (Supplemental Table S4) that fit the observed number of synonymous differences between a pair of species because there is a wide range of divergence times from the literature for each species. Here the number of synonymous differences equals 2 × divergence time × mutation rate. Second, using this divergence time, demography, and DFE inferred from Huber *et al.*(2017), or as described above for mice, we estimated the expected number nonsynonymous differences (*D*_*NE*_) according to Sawyer and Hartl eqn 13 (Sawyer and Hartl 1992; Boyko et al. 2008). Then, α is calculated as

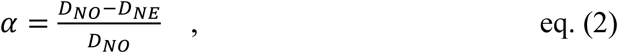

where *D*_*NO*_ is the observed number of nonsynonymous differences between species.

### DFE-alpha

Data files and the program v2.15 were downloaded from the following link: http://www.homepages.ed.ac.uk/pkeightl//dfe_alpha/download-dfe-alpha.html. Folded synonymous and nonsynonymous SFS were used as input in the inferences. The *est_alpha_omega* program was used to estimate the proportion of adaptive divergence.

### Composite likelihood approach for testing whether p^+^ and s^+^ differ across species

We first used *predicDiv* and *prfreq* to generate a look-up table for the expected number of nonsynonymous differences between species for a range of *s*^*+*^ (10^−5^-10^−2^) for each species. We focused on this range to capture strongly advantageous mutations. Given a demographic model, we again assume a substitution rate according to equation 13 in Sawyer and Hartl (1992). The contribution of the ancestral population to the divergence between two species is computed by numerically solving the diffusion equation using *predicDiv* and *prfreq* (Boyko et al. 2008). For each species, we then did a grid search of log_10_(*s*^*+*^) (−5 to -2) and *p*^*+*^(0-7.5%). We are interested in this range of strong *s*^*+*^ because weakly beneficial mutations still segregating as polymorphisms and are accounted for by the DFE fit to the SFS. We use a Poisson log-likelihood function to calculate the log-likelihood (LL) for each combination of *s*^*+*^ and *p*^*+*^. We find the MLE of *s*^*+*^ and *p*^*+*^ for each taxon under each demographic model that maximizes the LL and best fit the observed *D*_*N*_. This is the full model (H1) where each taxon (*i.e.* primates, rodent and *Drosophila*) is allowed to have its own *s*^*+*^ and *p*^*+*^. Our H0 hypothesis is the constrained model, where two or three taxa have the same *s*^*+*^ and *p*^*+*^. The LL of the constrained model is the sum of LL for each *s*^*+*^ and *p*^*+*^ for each taxon under comparison. We then found the MLE for the constrained model and calculate the likelihood ratio between H1 and H0.

### Composite likelihood approach for testing whether p^+^ and γ^+^ differ across species

Similarly, we used *predicDiv* and *prfreq* (Boyko et al. 2008) to generate a look-up table for the expected number of nonsynonymous differences for a range of ***γ***^*+*^ (1-10^3^) for each taxon under each demographic model. For each species, we performed a grid search of log_10_(***γ***^*+*^) (0-3) and *p*^*+*^(0-7.5%). Note, because effective population sizes differ over several orders of magnitude across our three taxa, we are searching across drastically different ranges of *s*^*+*^ as compared to our previous inference described above. We again use a Poisson log-likelihood function to calculate the LL for each combination of ***γ***^*+*^ and *p*^*+*^ and use a similar LRT framework as described above for *s*^*+*^ and *p*^*+*^.

### Composite parameters p^+^s^+^ and p^+^γ^+^

We examined two composite parameters, the product of *p*^*+*^ and *s*^*+*^ and the product of *p*^*+*^ and ***γ***^*+*^ for each taxon. Multiple combinations of parameters could give the same product *p*^*+*^*s*^*+*^ and *p*^*+*^***γ***^*+*^and distinct combinations could have different LL. Thus we found the values of *p*^*+*^ and *s*^*+*^ that gave the highest LL for each *p*^*+*^*s*^*+*^ value (and similarly each *p*^*+*^ and ***γ***^*+*^for each *p*^*+*^***γ***^*+*^). Values of log_10_(*p*^*+*^*s*^*+*^) and *p*^*+*^***γ***^*+*^were rounded to three digits prior to this comparison.

### Conditional log-likelihoods

To examine whether primates, rodents, and *Drosophila* could have the same *s*^*+*^ or *p*^*+*^, we examined the conditional log-likelihoods. To make the conditional log-likelihood curve for *p*^*+*^, for each *s*^*+*^ value, we find the *p*^*+*^ that maximizes the LL as well as the *p*^*+*^ values that have a LL within three LL units of this maximum LL for this *s*^*+*^ (*i.e. p*^*+*^| *s*^*+*^). To make the conditional log-likelihood curve for *s*^*+*^, for each *p*^*+*^ value, we look for the *s*^*+*^ that maximizes the log-likelihood as well as the *s*^*+*^ values that have a LL within three LL units of this maximum LL for this *p*^*+*^ (*i.e. s*^*+*^| *p*^*+*^). The same approach was used to construct the conditional log-likelihood curve for *p*^*+*^ and ***γ***^*+*^.

### Estimating α on the human lineage

Human-chimpanzee differences were polarized by the macaque sequence. Differences between human and chimpanzee were assigned to human lineage if the bases differ between human and macaque, but were the same between chimpanzee and macaque in the pairwise genome alignments. Differences that are in regions where the human and macaque sequences were un-alignable and sites that differ among human, chimp, and macaque cannot be polarized (3.8% total) and were filtered out. The total length of coding regions was scaled by this filter accordingly. A total of 30,530 synonymous substitutions and 20,013 nonsynonymous differences were observed on human lineage. We assume the human lineage and the outgroup lineage contribute equally to neutral *D*_*S*_, thus the total *D*_*S*_ between human and chimpanzee would be 61,060. We then adjusted divergence time to match predicted *D*_*S*_ to 61,060, and use this adjusted divergence time, DFE of deleterious mutations and demography of human to predict *D*_*N*_ as described in *Model-based estimates of α*. The human lineage α is then calculated using equation (2).

### Filtering to only include sites not affected by biased gene conversion or CpG hypermutation

Full polymorphism and substitutions datasets of primates and rodents were filtered to keep only strong to strong or weak to weak (SSWW) mutations because these changes were not affected by BGC or increased mutation rates due to deamination of methylated CpG sites. These changes include only A to T, T to A, C to G and G to C changes. These changes are only a small subset of all variable sites. The nonsynonymous and synonymous sequence lengths (*L*_*NS*_, *L*_*S*_) depend on the transition/transversion ratio and the CpG mutational bias. SSWW mutations are all transversions and do not included any CpG mutations, leading to a multiplier of *L*_*NS*_ = 5.21 x *L*_*S*_ in both primates and rodents. To compute this: 1) we used numbers of 0-, 2-, 3- and 4-fold sites in the human exome from Veeramah et al. (2014); 2) we consider all 2-fold sites to be nonsynonymous (because SSWW mutations are all transversions); and 3) we do not consider a mutational bias of CpG sites (because CpG sites are not included in the SSWW set). In addition, because SSWW mutations are only a small subset of all mutations, mutation rates need to be scaled down to the SSWW specific mutation rate. To estimate the demographic parameters and DFE for SSWW polymorphisms in rodents, first, we used the observed number of synonymous SSWW polymorphisms to estimate the mutation rate for mice SSWW mutations to be 5.99×10^−10^. Then, using the SFS for SSWW synonymous polymorphisms, we inferred that the ancestral population size in mice is approximately 246,256 which expanded 1.7-fold approximately 262,000 generations ago (Supplemental Table S3). Conditional on this demographic model, we estimated the DFE for new nonsynonymous SSWW mutations in mice. We assume that the DFE follows a gamma distribution and estimate its shape parameter α to be 0.21 and scale parameter beta to be 0.050 (Supplemental Table S3). These estimates are within the same magnitude of the estimates from the full dataset and a previous study (Huber et al. 2017). We re-estimated the mouse-rat divergence time that fits best with the observed SSWW *D*_*S*_. We then re-inferred *p*^*+*^, *s*^*+*^, and ***γ***^*+*^ as done previously, using the filtered data, new mutation rates, and new values of *L*_*NS*_.

## ACKNOWLEDGMENTS

We thank Lawrence Uricchio and David Enard for advice, discussions, and Tanya Phung, Jazlyn Mooney, Clare Marsden and Jesse Garcia for helpful comments on our manuscript. This work was supported by a Searle Scholars Fellowship and NIH Grant R35GM119856 (to K.E.L.) and foundation of Westlake University and NSFC 31900315 (to Y.Z.). We acknowledge support from a QCB Collaboratory Postdoctoral Fellowship to Y.Z. and the QCB Collaboratory Community directed by Matteo Pellegrini.

## Author contributions

K.E.L conceived of and supervised the study. Y.Z. carried out all analyses on positive selection and generated all figures. C.D.H carried demographic and gamma-DFE inference. R.W.D. processed mice raw data to genotypes. Y.Z., C.D.H., R.W.D and K.E.L. all participated in manuscript preparation.

## DISCLOSURE DECLARATION

The authors declare no competing interests.

## Notes

### Competing Interest Statement

The authors have declared no competing interest.

### Summary of Updates

We have more systematically explored the effect of the ancestral and outgroup population sizes on estimates of alpha. We also infer a composite parameter, p+s+, and show that this differs across species.

## REFERENCES

Andolfatto P. 2005. Adaptive evolution of non-coding DNA in Drosophila. Nature 437: 1149–1152.

Andolfatto P, Wong KM, Bachtrog D. 2011. Effective population size and the efficacy of selection on the X chromosomes of two closely related Drosophila species. Genome Biol Evol 3: 114–128.

Bolívar P, Mugal CF, Nater A, Ellegren H. 2015. Recombination rate variation modulates gene sequence evolution mainly via GC-biased gene conversion, not Hill–Robertson interference, in an avian system. Mol Biol Evol msv214.

Booker TR. 2019. Inferring parameters of the distribution of fitness effects of new mutations when beneficial mutations are strongly advantageous and rare. bioRxiv 855411.

Booker TR, Keightley PD. 2018. Understanding the factors that shape patterns of nucleotide diversity in the house mouse genome. Mol Biol Evol 35: 2971–2988.

Boyko AR, Williamson SH, Indap AR, Degenhardt JD, Hernandez RD, Lohmueller KE, Adams MD, Schmidt S, Sninsky JJ, Sunyaev SR, et al. 2008. Assessing the evolutionary impact of amino acid mutations in the human genome. PLoS Genet 4: e1000083.

Burgess R, Yang Z. 2008. Estimation of hominoid ancestral population sizes under bayesian coalescent models incorporating mutation rate variation and sequencing errors. Mol Biol Evol 25: 1979–1994.

Bustamante CD, Fledel-Alon A, Williamson S, Nielsen R, Hubisz MT, Glanowski S, Tanenbaum DM, White TJ, Sninsky JJ, Hernandez RD, et al. 2005. Natural selection on protein-coding genes in the human genome. Nature 437: 1153–1157.

Campos JL, Zhao L, Charlesworth B. 2017. Estimating the parameters of background selection and selective sweeps in Drosophila in the presence of gene conversion. Proc Natl Acad Sci 114: E4762–E4771.

Charlesworth J, Eyre-Walker A. 2008. The McDonald–Kreitman test and xlightly deleterious mutations. Mol Biol Evol 25: 1007–1015.

Chen F-C, Li W-H. 2001. Genomic divergences between humans and other hominoids and the effective population size of the common ancestor of humans and chimpanzees. Am J Hum Genet 68: 444–456.

Connallon T, Clark AG. 2015. The distribution of fitness effects in an uncertain world. Evol Int J Org Evol 69: 1610–1618.

The 1000 Genomes Project Consortium. 2012. An integrated map of genetic variation from 1,092 human genomes. Nature 491: 56–65.

Corbett-Detig RB, Hartl DL, Sackton TB. 2015. Natural selection constrains neutral diversity across a wide range of species. PLoS Biol 13: e1002112.

Corcoran P, Gossmann TI, Barton HJ, Slate J, Zeng K. 2017. Determinants of the efficacy of natural selection on coding and noncoding variability in two passerine species. Genome Biol Evol 9: 2987–3007.

Duncan BK, Miller JH. 1980. Mutagenic deamination of cytosine residues in DNA. Nature 287: 560.

Duret L, Galtier N. 2009. Biased Gene Conversion and the Evolution of Mammalian Genomic Landscapes. Annu Rev Genomics Hum Genet 10: 285–311.

Elyashiv E, Sattath S, Hu TT, Strutsovsky A, McVicker G, Andolfatto P, Coop G, Sella G. 2016. A genomic map of the effects of linked selection in Drosophila. PLOS Genet 12: e1006130.

Enard D, Messer PW, Petrov DA. 2014. Genome-wide signals of positive selection in human evolution. Genome Res 24: 885–895.

Eyre-Walker A. 2002. Changing effective population size and the McDonald-Kreitman test. Genetics 162: 2017–2024.

Eyre-Walker A, Keightley PD. 2009. Estimating the rate of adaptive molecular evolution in the presence of slightly deleterious mutations and population size change. Mol Biol Evol 26: 2097–2108.

Fay JC. 2011. Weighing the evidence for adaptation at the molecular level. Trends Genet 27: 343–349.

Fay JC, Wyckoff GJ, Wu CI. 2001. Positive and negative selection on the human genome. Genetics 158: 1227–1234.

Fay JC, Wyckoff GJ, Wu C-I. 2002. Testing the neutral theory of molecular evolution with genomic data from Drosophila. Nature 415: 1024–1026.

Fischer A, Wiebe V, Pääbo S, Przeworski M. 2004. Evidence for a complex demographic history of chimpanzees. Mol Biol Evol 21: 799–808.

Foxe JP, Dar V-N, Zheng H, Nordborg M, Gaut BS, Wright SI. 2008. Selection on amino acid substitutions in Arabidopsis. Mol Biol Evol 25: 1375–1383.

Galtier N. 2016. Adaptive Protein evolution in animals and the effective population size hypothesis. PLoS Genet 12: e1005774.

Gillespie JH. 2004. Why k=4Nus is silly. In The Evolution of Population Biology. (ed. RS Singh and MK Uyenoyama), pp. 178–192. Cambridge University Press, Cambridge, UK.

Gossmann TI, Keightley PD, Eyre-Walker A. 2012. The effect of variation in the effective population size on the rate of adaptive molecular evolution in eukaryotes. Genome Biol Evol 4: 658–667.

Gossmann TI, Song B-H, Windsor AJ, Mitchell-Olds T, Dixon CJ, Kapralov MV, Filatov DA, Eyre-Walker A. 2010. Genome wide analyses reveal little evidence for adaptive evolution in many plant species. Mol Biol Evol 27: 1822–1832.

Halligan DL, Kousathanas A, Ness RW, Harr B, Eöry L, Keane TM, Adams DJ, Keightley PD. 2013. Contributions of protein-coding and regulatory change to adaptive molecular evolution in murid rodents. PLoS Genet 9: e1003995.

Halligan DL, Oliver F, Eyre-Walker A, Harr B, Keightley PD. 2010. Evidence for pervasive adaptive protein evolution in wild mice. PLoS Genet 6: e1000825.

Hernandez RD, Hubisz MJ, Wheeler DA, Smith DG, Ferguson B, Rogers J, Nazareth L, Indap A, Bourquin T, McPherson J, et al. 2007. Demographic histories and patterns of linkage disequilibrium in Chinese and Indian rhesus macaques. Science 316: 240–243.

Hobolth A, Christensen OF, Mailund T, Schierup MH. 2007. Genomic relationships and speciation times of human, chimpanzee, and gorilla inferred from a coalescent hidden Markov model. PLoS Genet 3: e7.

Hu TT, Eisen MB, Thornton KR, Andolfatto P. 2013. A second-generation assembly of the Drosophila simulans genome provides new insights into patterns of lineage-specific divergence. Genome Res 23: 89–98.

Huber CD, Kim BY, Marsden CD, Lohmueller KE. 2017. Determining the factors driving selective effects of new nonsynonymous mutations. Proc Natl Acad Sci 114: 4465–4470.

Hvilsom C, Qian Y, Bataillon T, Li Y, Mailund T, Sallé B, Carlsen F, Li R, Zheng H, Jiang T, et al. 2012. Extensive X-linked adaptive evolution in central chimpanzees. Proc Natl Acad Sci 109: 2054–2059.

Keightley PD, Campos JL, Booker TR, Charlesworth B. 2016. Inferring the frequency spectrum of derived variants to quantify adaptive molecular evolution in protein- coding genes of Drosophila melanogaster. Genetics 203: 975–984.

Kimura M. 1962. On the probability of fixation of mutant genes in a population. Genetics 47: 713–719.

Kimura M. 1983. The Neutral Theory of Molecular Evolution. Cambridge University Press, Cambridge.

Lachance J, Tishkoff SA. 2014. Biased gene conversion skews allele frequencies in human populations, increasing the disease burden of recessive alleles. Am J Hum Genet 95: 408–420.

Lack JB, Cardeno CM, Crepeau MW, Taylor W, Corbett-Detig RB, Stevens KA, Langley CH, Pool JE. 2015. The Drosophila Genome Nexus: A population genomic resource of 623 *Drosophila melanogaster* genomes, including 197 from a single ancestral range population. Genetics 199: 1229–1241.

Li H, Durbin R. 2009. Fast and accurate short read alignment with Burrows-Wheeler transform. Bioinforma Oxf Engl 25: 1754–1760.

Lourenço JM, Glémin S, Galtier N. 2013. The rate of molecular adaptation in a changing environment. Mol Biol Evol 30: 1292–1301.

Lunter G, Goodson M. 2011. Stampy: a statistical algorithm for sensitive and fast mapping of Illumina sequence reads. Genome Res 21: 936–939.

McDonald JH, Kreitman M. 1991. Adaptive protein evolution at the Adh locus in Drosophila. Nature 351: 652–654.

McKenna A, Hanna M, Banks E, Sivachenko A, Cibulskis K, Kernytsky A, Garimella K, Altshuler D, Gabriel S, Daly M, et al. 2010. The Genome Analysis Toolkit: a MapReduce framework for analyzing next-generation DNA sequencing data. Genome Res 20: 1297–1303.

McVicker G, Gordon D, Davis C, Green P. 2009. Widespread genomic signatures of natural selection in hominid evolution. PLoS Genet 5: e1000471.

Messer PW, Petrov DA. 2013. Frequent adaptation and the McDonald–Kreitman test. Proc Natl Acad Sci 110: 8615–8620.

Nam K, Munch K, Mailund T, Nater A, Greminger MP, Krützen M, Marquès-Bonet T, Schierup MH. 2017. Evidence that the rate of strong selective sweeps increases with population size in the great apes. Proc Natl Acad Sci U S A 114: 1613–1618.

Ness RW, Zhang Y-H, Cong L, Wang Y, Zhang J-X, Keightley PD. 2012. Nuclear gene variation in wild brown rats. G3 Genes Genomes Genet 2: 1661–1664.

Ohta T. 1973. Slightly deleterious mutant substitutions in evolution. Nature 246: 96–98.

Ohta T. 1992. The nearly neutral theory of molecular evolution. Annu Rev Ecol Syst 23: 263–286.

Orr HA. 1998. The population genetics of adaptation: The distribution of factors fixed during adaptive evolution. Evolution 52: 935–949.

Phifer-Rixey M, Bonhomme F, Boursot P, Churchill GA, Piálek J, Tucker PK, Nachman MW. 2012. Adaptive evolution and effective population size in wild house mice. Mol Biol Evol 29: 2949–2955.

Prado-Martinez J, Sudmant PH, Kidd JM, Li H, Kelley JL, Lorente-Galdos B, Veeramah KR, Woerner AE, O’Connor TD, Santpere G, et al. 2013. Great ape genetic diversity and population history. Nature 499: 471–475.

Pritchard JK, Pickrell JK, Coop G. 2010. The genetics of human adaptation: hard sweeps, soft sweeps, and polygenic adaptation. Curr Biol 20: R208–R215.

Robinson MC, Stone EA, Singh ND. 2014. Population genomic analysis reveals no evidence for GC-biased gene conversion in *Drosophila melanogaster*. Mol Biol Evol 31: 425–433.

Romiguier J, Gayral P, Ballenghien M, Bernard A, Cahais V, Chenuil A, Chiari Y, Dernat R, Duret L, Faivre N, et al. 2014. Comparative population genomics in animals uncovers the determinants of genetic diversity. Nature 515: 261–263.

Rousselle M, Mollion M, Nabholz B, Bataillon T, Galtier N. 2018. Overestimation of the adaptive substitution rate in fluctuating populations. Biol Lett 14: 20180055.

Rousselle M, Simion P, Tilak MK, Figuet E, Nabholz B, Galtier N. 2019. Is adaptation limited by mutation? A timescale-dependent effect of genetic diversity on the adaptive substitution rate in animals. bioRxiv 643619.

Sawyer SA, Hartl DL. 1992. Population genetics of polymorphism and divergence. Genetics 132: 1161–1176.

Schneider A, Charlesworth B, Eyre-Walker A, Keightley PD. 2011. A method for inferring the rate of occurrence and fitness effects of advantageous mutations. Genetics 189: 1427–1437.

Schrago CG. 2014. The effective population sizes of the anthropoid ancestors of the human– chimpanzee lineage provide insights on the historical biogeography of the great apes. Mol Biol Evol 31: 37–47.

Sella G, Petrov DA, Przeworski M, Andolfatto P. 2009. Pervasive natural selection in the Drosophila genome? PLoS Genet 5: e1000495.

Smith NGC, Eyre-Walker A. 2002. Adaptive protein evolution in Drosophila. Nature 415: 1022–1024.

Strasburg JL, Kane NC, Raduski AR, Bonin A, Michelmore R, Rieseberg LH. 2011. Effective population size is positively correlated with levels of adaptive divergence among annual sunflowers. Mol Biol Evol 28: 1569–1580.

Stumpf MPH, Thorne T, Silva E de, Stewart R, An HJ, Lappe M, Wiuf C. 2008. Estimating the size of the human interactome. Proc Natl Acad Sci 105: 6959–6964.

Sved J, Bird A. 1990. The expected equilibrium of the CpG dinucleotide in vertebrate genomes under a mutation model. Proc Natl Acad Sci U S A 87: 4692–4696.

Tataru P, Mollion M, Glémin S, Bataillon T. 2017. Inference of distribution of fitness effects and proportion of adaptive substitutions from polymorphism data. Genetics 207: 1103–1119.

Torgerson DG, Boyko AR, Hernandez RD, Indap A, Hu X, White TJ, Sninsky JJ, Cargill M, Adams MD, Bustamante CD, et al. 2009. Evolutionary processes acting on candidate cis-regulatory regions in humans inferred from patterns of polymorphism and divergence. PLoS Genet 5: e1000592.

Uricchio LH, Petrov DA, Enard D. 2019. Exploiting selection at linked sites to infer the rate and strength of adaptation. Nat Ecol Evol 3: 977–984.

Valentine JW, Collins AG, Meyer CP. 1994. Morphological complexity increase in metazoans. Paleobiology 20: 131–142.

Veeramah KR, Gutenkunst RN, Woerner AE, Watkins JC, Hammer MF. 2014. Evidence for increased levels of positive and negative selection on the X chromosome versus autosomes in humans. Mol Biol Evol 31: 2267–2282.

Wall JD. 2013. Great Ape Genomics. ILAR J 54: 82–90.

Xue C, Raveendran M, Harris RA, Fawcett GL, Liu X, White S, Dahdouli M, Rio Deiros D, Below JE, Salerno W, et al. 2016. The population genomics of rhesus macaques (Macaca mulatta) based on whole-genome sequences. Genome Res 26: 1651–1662.

Yang H, Ding Y, Hutchins LN, Szatkiewicz J, Bell TA, Paigen BJ, Graber JH, de Villena FP-M, Churchill GA. 2009. A customized and versatile high-density genotyping array for the mouse. Nat Methods 6: 663–666.

