## Supplementary Information for "Greater strength of selection and higher proportion of beneficial amino acid changing mutations in humans compared to mice and *Drosophila melanogaster*"

### Table of Contents


- Supplemental Text
- Supplemental Figures S1-S8
- Supplemental Tables S1-S6

### Supplemental Text

#### *Demographic and DFE inferences for mice*

We used methods established in Huber et al. (2017) to infer demography and the DFE of neutral and deleterious mutations from mouse polymorphism data. In short, we first used the synonymous SFS to infer demographic parameters for the Simple model using ∂a∂i (Sawyer and Hartl 1992; Gutenkunst et al. 2009). We infer that the ancestral population size is approximately 206,500 which expanded 2.4-fold 293,000 generations ago (Supplemental Table S3). Conditional on this demographic model, we estimated the DFE for new nonsynonymous mutations in mice. We assume that the DFE follows a gamma-distribution and estimate its shape parameter $\alpha$ to be 0.21 and scale parameter to be 0.083 (Supplemental Table S3). These estimates are within the same magnitude of previous estimates from Huber *et al.* (2017), which used a much smaller dataset (<0.1% of the total sites used in our study).

#### Details on the analysis of SSWW sites

To test whether BGC and hypermutable CpG sites drive the observed pattern of positive selection across species, we filtered human and mouse data to keep only strong to strong or weak to weak mutations (herein called SSWW mutations), which are not affected by BGC and are not CpG changes. For humans, the filtered SSWW polymorphisms have a similar SFS as the full dataset (Supplemental Fig. S5A). Thus we use the demographic and DFE parameters estimated from the full data. We used the observed number of synonymous SSWW polymorphisms to estimate the mutation rate of human SSWW mutations to be $3.14\times10^{-9}$, which is comparable to previous estimates (Kong et al. 2012; Lachance and Tishkoff 2014). Following the method used for the full data, we re-estimated the human-chimpanzee divergence time that fits best to the observed SSWW $D_S$. We then predict the SSWW $D_N$ using this newly estimated divergence time, DFE, and demographic models (Supplemental Table S1). We estimated that under the Simple demographic model (*Nout=Nanc=Nanc.in*) and the Complex demographic model (*Nout≠Nanc≠Nanc.in*), approximately 13.1% or approximately 26.3% of the observed SSWW $D_N$ in humans using chimpanzee as outgroup was driven by positive selection, respectively (Table 1). These estimates of α from SSWW sites are slightly elevated but are comparable to the estimates from the full dataset (Table 1).

For mouse, however, the SFS of SSWW polymorphism has a very different shape compared to the SFS from the full dataset (Supplemental Fig. S5B). Thus, we re-estimated the demographic and DFE parameters for the mouse SSWW mutations (Supplemental Table S3; see Methods). We then estimated that under the Simple demographic model and the Complex demographic model, approximately 19.5% and 9.7% of the observed SSWW $D_N$ in mice was driven by positive selection, respectively (Table 1). These estimates of proportion of nonsynonymous substitutions fixed by positive selection from SSWW mutations are much lower than those estimated from the full dataset (Table 1). This suggests that biased gene conversion and CpG mutational processes may account for some of the nonsynonymous substitutions between mouse and rat.

***Coalescent simulations to compare human polymorphism under Simple and Complex demographic models***

To evaluate whether patterns of neutral polymorphism would be predicted to be different under the human Simple and Complex demographic models, we conducted coalescent simulations under these models using *ms* (Hudson 2002). Specifically, we simulated 1000 replicates for each scenario and calculated the mean number of synonymous segregating sites across replicates. Both models showed similar numbers of neutral segregating sites (34075 for the Simple Model and 33810 for the Complex model), suggesting that using population sizes more appropriate for the outgroup population will not affect polymorphism data in the ingroup sample.

Supplemental Fig. S1
Complex demographic model used for divergence between human (ingroup) and macaque (outgroup). In this demographic model, at the timing of the human-chimp split approximately 105,000 generations ago (2.6 Myr assuming 25 years/generation), the human population changes to that of the human-chimp ancestral size of 60,000 (see Results). *Nout* is set to 73,000 (Hernandez et al. 2011).

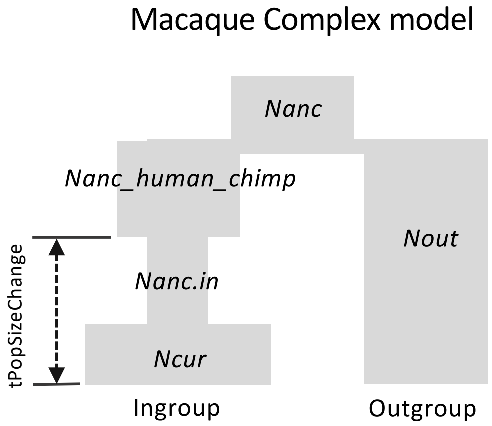

Supplemental Fig. S2

Observed and predicted number of nonsynonymous divergent sites between each of our three focal species and their outgroups. (A) Predictions using MLEs of $p^+$ and $s^+$. (B) Predictions using MLEs of $p^+$ and $\boldsymbol{\gamma}^+$. Predicted numbers are from two models: the full model, H1, where each species has its own positive selection parameters and the constrained model, H0, where the positive selection parameters are constrained to be the same across all three taxa.

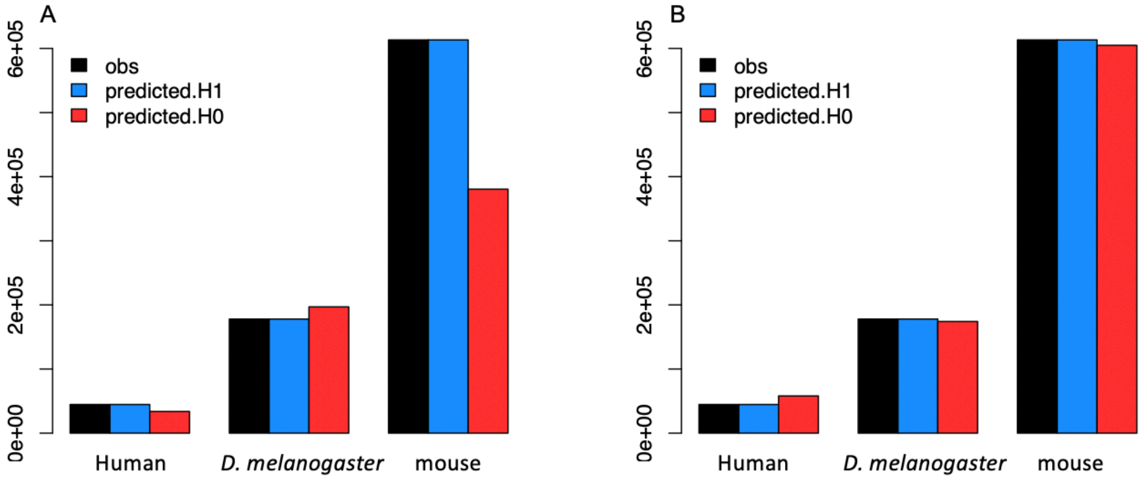

Supplemental Fig. S3

Conditional log-likelihood (LL) surfaces. (A) Maximizing $p^+$ given particular values of $\boldsymbol{\gamma}^+$ and (B) maximizing $\boldsymbol{\gamma}^+$ given particular values of $p^+$. Only grid points within 3 LL units of the MLEs of for each parameter for each species are shown. Light blue denotes human, pink denotes *D. melanogaster*, and light green denotes mouse.

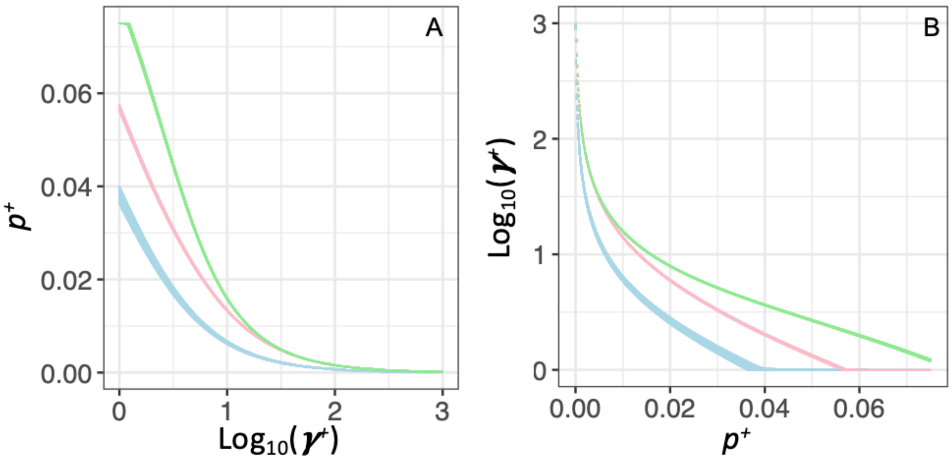

Supplemental Fig. S4

The composite parameter $p^+\boldsymbol{\gamma}^+$ across species. Log-likelihood surfaces for $p^+\boldsymbol{\gamma}^+$ in the three different taxa. Red denotes the inference for *D. melanogaster*, green denotes the inference for mouse, blue denotes the inference for humans using the chimpanzee as the outgroup. Lighter colors denote the Simple model. Darker colors denote the Complex model that better models the ancestral demography and population size of the outgroup.

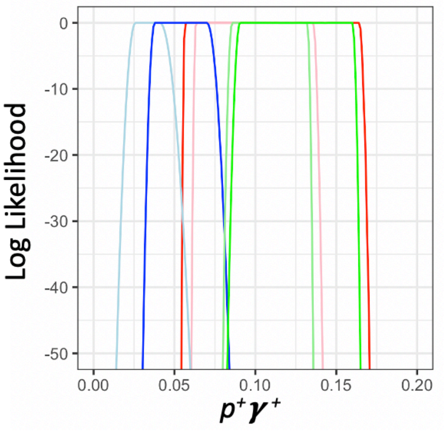

Supplemental Fig. S5
Effects of biased gene conversion on the folded SFS for (A) humans and (B) mice. Full denotes the data without any filtering for biased gene conversion. SSWW denotes the SFS for strong to strong or weak to weak substitutions only. S denotes synonymous SNPs. NS denotes nonsynonymous SNPs.

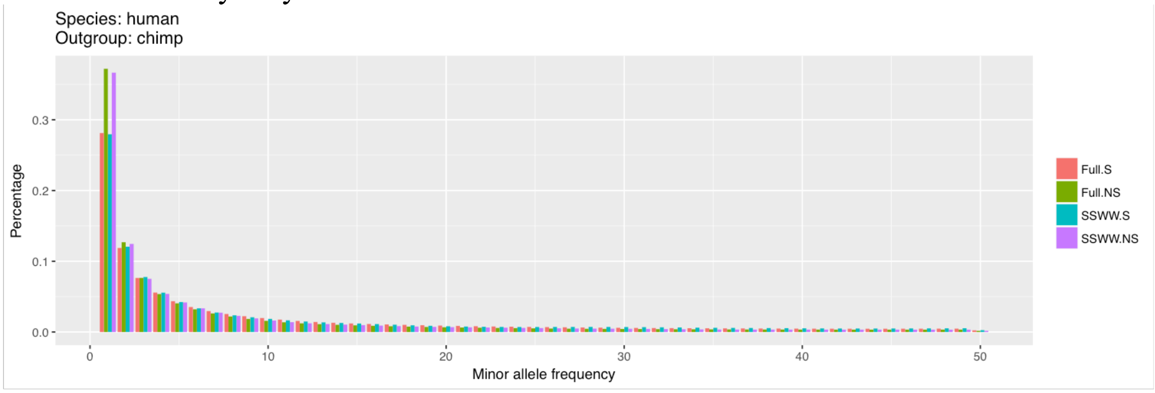


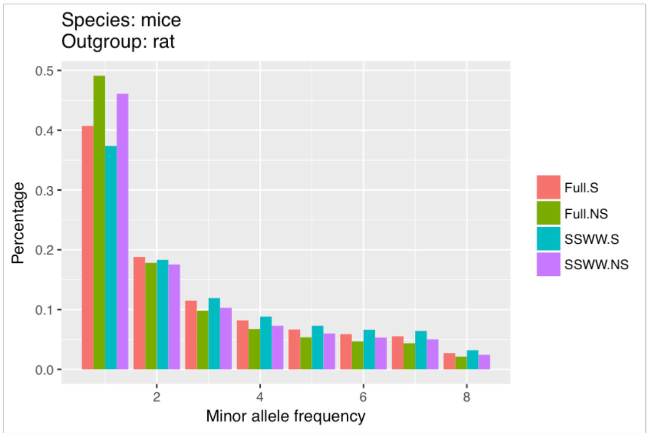

Supplemental Fig. S6

Log-likelihood surfaces for sites unaffected by biased gene conversion (SSWW sites in mammals). (A-C) show the log-likelihood surfaces for $p^+$ and $s^+$ for different taxa. (D) shows the constrained model, H0, where $p^+$ and $s^+$ are constrained to be the same across all three taxa. Log-Likelihoods are calculated using grid search method of $\log_{10}(s^+)$ in the range of -5 to -2 and $p^+$ in the range of 0-7.5%. Blue denotes human, red denotes *D. melanogaster*, and green denotes mouse. The large points represent the MLE for each species. The black cross in panel D represents the MLE of the constrained model, and the lighter colors show grid points within 3 LL units of each MLE. The Complex model is used for each species and we use chimpanzee as the outgroup for humans. (E) shows the conditional log-likelihood surface maximizing $p^+$ given particular values of $s^+$ and (F) shows the conditional log-likelihood surface maximizing $s^+$ given particular values of $p^+$. In panels E-F, only grid points within 3 LL units of the MLEs of for each parameter for each species are shown.

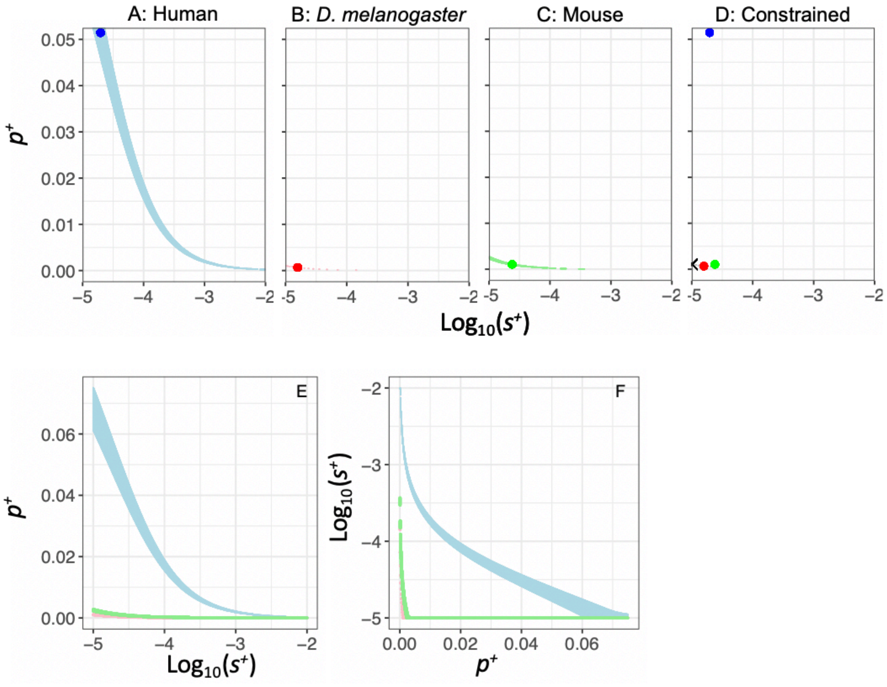

Supplemental Fig. S7

Log-likelihood (LL) surfaces for sites unaffected by biased gene conversion (SSWW sites in mammals) for $p^+$ and $\boldsymbol{\gamma}^+$. (A-C) show the log-likelihood surfaces for $p^+$ and $\boldsymbol{\gamma}^+$ for different taxa. (D) shows the constrained model, H0, where $p^+$ and $\boldsymbol{\gamma}^+$ are constrained to be the same across all three taxa. Log-Likelihoods are calculated using grid search method of $\log_{10}(\boldsymbol{\gamma}^+)$ in the range of 0 to 3 and $p^+$ in the range of 0-7.5%. Blue denotes human, red denotes *D. melanogaster*, and green denotes mouse. The large points represent the MLE for each species. The black cross in panel D represents the MLE of the constrained model, and the lighter colors show grid points within 3 LL units of each MLE. The Complex model is used for each species and we use chimpanzee as the outgroup for humans. (E) shows the conditional log-likelihood surface maximizing $p^+$ given particular values of $\boldsymbol{\gamma}^+$ and (F) shows the conditional log-likelihood surface maximizing $\boldsymbol{\gamma}^+$ given particular values of $p^+$. In panels E-F, only grid points within 3 LL units of the MLEs of for each parameter for each species are shown.

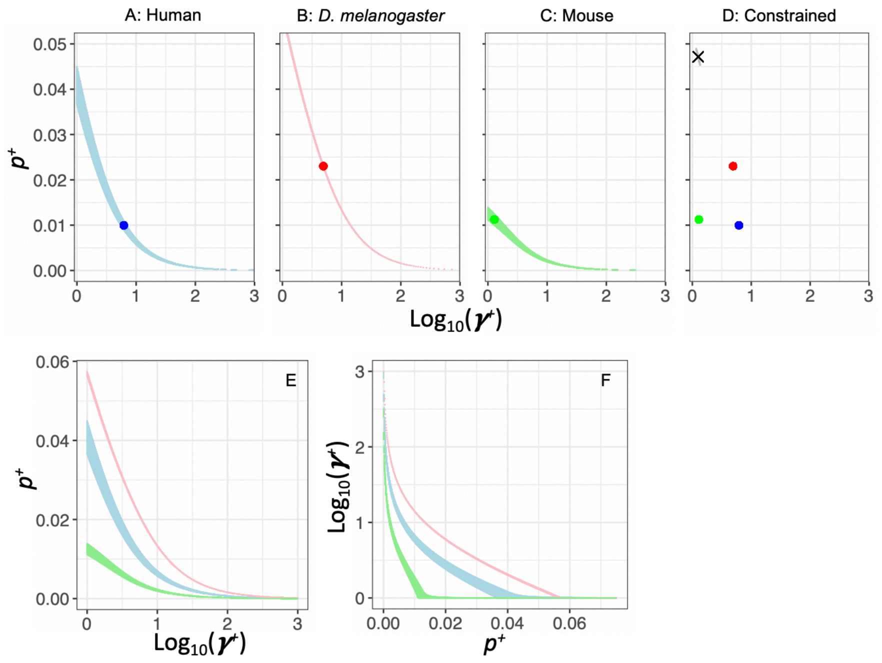

Supplemental Fig. S8

The composite parameters $p^+s^+$ and $p^+\boldsymbol{\gamma}^+$ across taxa using SSWW sites that are not influenced by BGC or hypermutable CpG sites. A. Log-likelihood surfaces for $p^+s^+$ in the three different taxa. B. Log-likelihood surfaces for $p^+\boldsymbol{\gamma}^+$ in the three different taxa. Red denotes the inference for *D. melanogaster*, green denotes the inference for mouse, blue denotes the inference for humans using the chimpanzee as the outgroup. Lighter colors denote the Simple model. Darker colors denote the Complex model that better models the ancestral demography and population size of the outgroup.

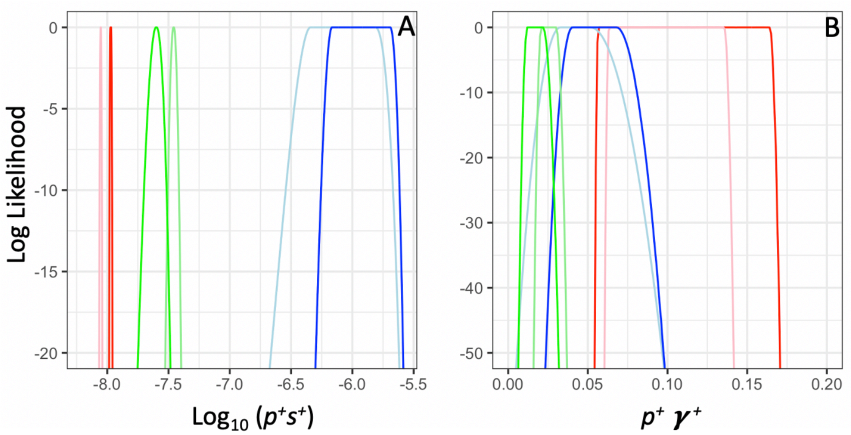

Supplemental Table S1. Observed and predicted counts of polymorphism and divergence.

| species | outgroup | demographic model | dataset | Ds | Dn | Ps | Pn |
|---|---|---|---|---|---|---|---|
| human | chimpanzee | observed | full | 65,021 | 44,665 | 33,684 | 32,520 |
| | | predicted:Simple Model | | 65,021 | 39,925 | | |
| | | predicted:Complex Model | | 65,021 | 33,681 | | |
| | macaque | observed | | 323,581 | 182,532 | 34,113 | 32,798 |
| | | predicted:Simple Model | | 323,581 | 196,401 | | |
| | | predicted:Complex Model | | 323,581 | 135,101 | | |
| | chimpanzee | observed | SSWW | 5,077 | 8,084 | 2,344 | 5,116 |
| | | predicted:Simple Model | | 5,077 | 7,026 | | |
| | | predicted:Complex Model | | 5,077 | 5,961 | | |
| *D. melanogaster* | *D. simulans* | observed | full | 404,537 | 177,936 | 466,188 | 231,267 |
| | | predicted:Simple Model | | 404,537 | 83,193 | | |
| | | predicted:Complex Model | | 404,537 | 70,660 | | |
| mice | rat | observed | full | 1,133,269 | 613,281 | 181,039 | 73,591 |
| | | predicted:Simple Model | | 1,133,269 | 334,398 | | |
| | | predicted:Complex Model | | 1,133,269 | 360,664 | | |
| | | observed | SSWW | 58,882 | 51,462 | 10,713 | 10,662 |
| | | predicted:Simple Model | | 58,882 | 41,422 | | |
| | | predicted:Complex Model | | 58,882 | 46,471 | | |

Predicted counts of $D_n$ come from the gamma-DFE with only neutral and deleterious mutations. The differences between observed and predicted $D_n$ are attributed to positive selection and are used to estimate $\alpha$.

Supplemental Table S2. Effect of slightly deleterious mutations on α.

| | | | MAF* | | | | |
|---|---|---|---|---|---|---|---|
| **Species** | **Outgroup** | **Sample Size** | **0** | **>5%** | **>10%** | **>20%** | **>30%** |
| Human | Chimpanzee | 100 | -0.41 | -0.09 | -0.04 | -0.01 | 0.00 |
| Human | Macaque | 100 | -0.70 | -0.33 | -0.26 | -0.22 | -0.21 |
| *D. melanogaster* | *D. simulans* | 100 | -0.13 | 0.39 | 0.44 | 0.49 | 0.51 |
| Mouse | Rat | 16 | 0.25 | na | 0.36 | 0.40 | 0.40 |

*MAF=0 includes SNPs at all frequencies when computing α. MAF>5% includes only SNPs at frequencies larger than 5%, etc.

Supplemental Table S3. Demographic and DFE parameters estimated from polymorphism data.

| parameter (unit) | human | Drosophila | mice | human SSWW | mice SSWW |
|---|---|---|---|---|---|
| sample size | 100 | 100 | 16 | 100 | 16 |
| total sites | 19089129 | 15819843 | 26642307 | 19089129 | 26642307 |
| Nanc | 7067 | 2790000 | 206519 | 7067 | 246256 |
| Ncurr/Nanc | 2.34 | 2.73 | 2.37 | 2.34 | 1.71 |
| tau (2Nanc) | 0.43 | 0.09 | 0.71 | 0.43 | 0.53 |
| DFE: alpha | 0.19 | 0.35 | 0.21 | 0.19 | 0.21 |
| DFE: beta (s) | 0.074 | 0.00038 | 0.083 | 0.074 | 0.050 |
| mu | 2.5E-08 | 1.5E-09 | 5.4E-09 | 3.14E-09 | 5.99E-10 |
| NS/S length ratio | 2.31 | 2.85 | 2.31 | 5.21 | 5.21 |

Nanc: ancestral population size. Ncurr: current population size. Tau: is the time at which the population changed in size. NS:nonsynonymous. S: synonymous. mu: mutation rate per site per generation.

Supplemental Table S4. Parameters of the Simple demographic model and the Complex demographic model for each species

| species | human | | | | | | *D. melanogaster* | | mice | | | |
|---|---|---|---|---|---|---|---|---|---|---|---|---|
| demographic model | Simple Model | Complex Model | Simple Model | Complex Model | Simple Model | Complex Model | Simple Model | Complex Model | Simple Model | Complex Model | Simple Model | Complex Model |
| outgroup | chimpanzee | chimpanzee | macaque | macaque | chimpanzee | chimpanzee | *D. simulans* | *D. simulans* | rat | rat | rat | rat |
| dataset | full | full | full | full | SSWW | SSWW | full | full | full | full | SSWW | SSWW |
| sample size | 100 | 100 | 100 | 100 | 100 | 100 | 100 | 100 | 16 | 16 | 16 | 16 |
| syn theta | 9538.37 | 9538.37 | 9620.51 | 9620.51 | 656.78 | 656.78 | 187784.40 | 187784.40 | 85008.11 | 85008.11 | 4329.45 | 4329.45 |
| tdiv | 6.39 | 3.19 | 33.21 | 30.73 | 7.30 | 4.10 | 1.79 | 0.65 | 12.91 | 13.13 | 13.02 | 13.40 |
| tau | 0.18 | 0.18 | 0.18 | 0.18 | 0.18 | 0.18 | 0.03 | 0.03 | 0.30 | 0.30 | 0.31 | 0.31 |
| omega | 0.43 | 0.43 | 0.43 | 0.43 | 0.43 | 0.43 | 0.37 | 0.37 | 0.42 | 0.42 | 0.58 | 0.58 |
| omega_outgroup | 0.43 | 1.81 | 0.43 | 4.41 | 0.43 | 1.81 | 0.37 | 1.50 | 0.42 | 0.20 | 0.58 | 0.20 |
| omega_ancestral | 0.43 | 3.63 | 0.43 | 2.90 | 0.43 | 3.63 | 0.37 | 1.50 | 0.42 | 0.20 | 0.58 | 0.20 |

All scaled to current ingroup population size (Ncurr in Supplemental Table S3).
tdiv: divergence time of ingroup and outgroup species;
tau: time from present to the first change in ingroup Ne;
omega: ancestral ingroup population size;
omega_outgroup: outgroup long-term effective population size;
omega_ancetral: ancestral population size before ingroup and outgroup divergence

Only for the Complex model for human using the macaque as the outgroup, we have two additional parameters: omega2 and tPopSizeChange to model the change of human ingroup population size to human-chimpanzee ancestral population size (omega2=3.63) at human-chimpanzee divergence time (tPopSizeChange=3.19).

Supplemental Table S5. Effective population sizes of chimpanzee, macaque, and ancestral primate populations from the literature.

| | Ne | Reference |
|---|---|---|
| chimpanzee | 30900–61800 | Prado-Martinez et al 2013 |
| | 25000-35000 | Fisher et al 2004 |
| | 18300 | Hvilsom et al 2014 |
| | ~30000 | Hvilsom et al 2012 |
| human-chimpanzee ancestor | 52,000–96,000 | Chen and Li 2001 |
| | 12,000–21,000 | Yang et al 2002 |
| | 65000 | Hobolth et al 2007 |
| | 47,000 | Hobolth et al 2011 |
| | 35,000-65,000 | Ruvolo et al 1997 |
| | 99 (95–102) x 1000 | Burgess et al 2008 |
| | 50000, 63000 | Prado-Martinez et al 2013 |
| | 33000 | Hara et al 2012 |
| | 27716-41263 | Schrago et al 2013 |
| | ~50000 | Wall 2003 |
| | 47500 | Schrago 2014 |
| macaque | 73000 (ancestral) | Hernandez et al 2007 |
| | 52350, 61800 (Indian rhesus) | Xue et al 2016 |
| | 71200, 82080 (Chinese rhesus) | Xue et al 2016 |
| human-macaque ancestor | 48000 | McVicker et al 2009 |

Supplemental Table S6. Comparison of the beneficial selection coefficients and proportion of new beneficial mutations across all three taxa.

A. Test whether $s^+$ and $p^+$ differ across taxa using the full datasets

| 1 | Hypothesis | species | outgroup | demorgraphic model | p+ | log10(s+) | LL | abbr. |
|---|---|---|---|---|---|---|---|---|
| | Full model (H1) | **Human** | **chimpanzee** | **Complex** | **1.55E-02** | **-3.949** | **-6.27** | **human3** |
| | | Human | Macaque | Complex | 7.20E-03 | -4.422 | -6.98 | human3mac |
| | | Human | chimpanzee | Simple | 2.39E-02 | -4.429 | -6.27 | human2 |
| | | ***D.melanogaster*** | ***D.simulans*** | **Complex** | **6.75E-04** | **-4.801** | **-6.96** | **fly3** |
| | | *D.melanogaster* | *D.simulans* | Simple | 6.00E-04 | -4.831 | -6.96 | fly2 |
| | | **Mouse** | **Rat** | **Complex** | **1.02E-02** | **-4.797** | **-7.58** | **mice3** |
| | | Mouse | Rat | Simple | 1.21E-02 | -4.954 | -7.58 | mice2 |

| 2 | | models comparison | sum of LL | p+ | log10(s+) | constrained LL | likelihood ratio | p-value |
|---|---|---|---|---|---|---|---|---|
| | Constrained (H0): same s+, p+ | **human3=fly3=mice3** | **-20.82** | **7.50E-05** | **-3.772** | **-62507.88** | **124974.12** | <1E-16 |
| | | **human3=fly3** | **-13.24** | **1.05E-03** | **-4.995** | **-1576.27** | **3126.07** | <1E-16 |
| | | human3=fly2 | -13.24 | 8.25E-04 | -4.969 | -1588.05 | 3149.63 | <1E-16 |
| | | human2=fly3 | -13.24 | 1.05E-03 | -4.995 | -268.00 | 509.52 | <1E-16 |
| | | human2=fly2 | -13.24 | 8.25E-04 | -4.970 | -271.30 | 516.12 | <1E-16 |
| | | human3mac=fly3 | -13.94 | 1.05E-03 | -4.992 | -6835.68 | 13643.47 | <1E-16 |
| | | human3mac=fly2 | -13.94 | 9.00E-04 | -5.000 | -6943.80 | 13859.73 | <1E-16 |
| | | **human3=mice3** | **-13.85** | **1.64E-02** | -5.000 | **-852.13** | **1676.56** | <1E-16 |
| | | human3=mice2 | -13.85 | 1.35E-02 | -5.000 | -968.96 | 1910.21 | <1E-16 |
| | | human2=mice3 | -13.85 | 1.64E-02 | -5.000 | -92.47 | 157.22 | <1E-16 |
| | | human2=mice2 | -13.85 | 1.34E-02 | -5.000 | -117.56 | 207.41 | <1E-16 |
| | | human3mac=mice3 | -14.56 | 1.68E-02 | -5.000 | -840.342 | 1651.57 | <1E-16 |
| | | human3mac=mice2 | -14.56 | 1.38E-02 | -5.000 | -1487.82 | 2946.52 | <1E-16 |
| | | **fly3=mice3** | **-14.55** | **7.50E-05** | **-3.772** | **-60903.15** | **121777.21** | <1E-16 |
| | | fly3=mice2 | -14.55 | 7.50E-05 | -3.735 | -73462.87 | 146896.65 | <1E-16 |
| | | fly2=mice3 | -14.55 | 7.50E-05 | -3.852 | -62865.88 | 125702.67 | <1E-16 |
| | | fly2=mice2 | -14.55 | 7.50E-05 | -3.812 | -76531.50 | 153033.91 | <1E-16 |

B. Test whether $\gamma^+$ and $p^+$ differ across taxa using the full datasets

| 1 | model | species | outgroup | demorgraphic model | $p^+$ | $\gamma^+$ | log-likelihood | abbr. |
|---|---|---|---|---|---|---|---|---|
| | Full model (H1) | **Human** | **chimpanzee** | **Complex** | **0.01** | **6.05** | **-6.27** | **human3** |
| | | Human | Macaque | Complex | 0.01 | 1.17 | -6.98 | human3mac |
| | | Human | chimpanzee | Simple | 0.02 | 1.65 | -6.27 | human2 |
| | | ***D.melanogaster*** | ***D.simulans*** | **Complex** | **0.02** | **4.92** | **-6.96** | **fly3** |
| | | *D.melanogaster* | *D.simulans* | Simple | 0.04 | 2.79 | -6.96 | fly2 |
| | | **Mouse** | **Rat** | **Complex** | **0.04** | **3.76** | **-7.58** | **mice3** |
| | | Mouse | Rat | Simple | 0.05 | 2.15 | -7.58 | mice2 |

| 2 | | models comparison | sum of LL | $p^+$ | $\gamma^+$ | constrained LL | likelihood ratio | p-value |
|---|---|---|---|---|---|---|---|---|
| | Constrained (H0): same gamma, p+ | **human3=fly3=mice3** | **-20.82** | **0.00** | **61.66** | **-1791.51** | **3541.39** | <1E-16 |
| | | **human3=fly3** | **-13.24** | **0.06** | **1.00** | **-292.10** | **557.73** | <1E-16 |
| | | human3=fly2 | -13.24 | 0.06 | 1.00 | -498.49 | 970.50 | <1E-16 |
| | | human2=fly3 | -13.24 | 0.06 | 1.00 | -319.50 | 612.54 | <1E-16 |
| | | human2=fly2 | -13.24 | 0.06 | 1.00 | -455.30 | 884.14 | <1E-16 |
| | | human3mac=fly3 | -13.94 | 0.02 | 1.00 | -26283.69 | 52539.50 | <1E-16 |
| | | human3mac=fly2 | -13.94 | 0.02 | 1.00 | -22821.27 | 45614.66 | <1E-16 |
| | | **human3=mice3** | **-13.85** | **0.08** | **1.00** | **-1309.62** | **2591.53** | <1E-16 |
| | | human3=mice2 | -13.85 | 0.00 | 845.28 | -859.97 | 1692.23 | <1E-16 |
| | | human2=mice3 | -13.85 | 0.08 | 1.10 | -936.87 | 1846.03 | <1E-16 |
| | | human2=mice2 | -13.85 | 0.08 | 1.09 | -895.77 | 1763.84 | <1E-16 |
| | | human3mac=mice3 | -14.56 | 0.02 | 1.00 | -44692.64 | 89356.16 | <1E-16 |
| | | human3mac=mice2 | -13.85 | 0.02 | 1.00 | -54279.71 | 108531.71 | <1E-16 |
| | | **fly3=mice3** | **-14.55** | **0.00** | **58.48** | **-14.55** | **0.00** | **1** |
| | | fly3=mice2 | -14.55 | 0.01 | 9.08 | -14.56 | 0.02 | 0.99 |
| | | fly2=mice3 | -14.55 | 0.00 | 993.12 | -424.24 | 819.39 | <1E-16 |
| | | fly2=mice2 | -14.55 | 0.01 | 12.76 | -14.56 | 0.02 | 0.99 |

C. Test whether $s^+$ and $p^+$ differ across taxa using only SSWW changes for human and mouse

| [1] Hypothesis | species | outgroup | demorgraphic model | p+ | log10(s+) | LL | abbr. | |
|---|---|---|---|---|---|---|---|---|
| Full model (H1) | **Human** | **chimpanzee** | **Complex** | **5.15E-02** | **-4.706** | **-5.42** | **human3bgc** | |
| | Human | chimpanzee | Simple | 3.26E-02 | -4.500 | -5.42 | human2bgc | |
| | ***D.melanogaster*** | ***D.simulans*** | **Complex** | **6.75E-04** | **-4.801** | **-6.96** | **fly3** | |
| | *D.melanogaster* | *D.simulans* | Simple | 6.00E-04 | -4.831 | -6.96 | fly2 | |
| | **Mouse** | **Rat** | **Complex** | **1.05E-03** | **-4.620** | **-6.34** | **mice3bgc** | |
| | Mouse | Rat | Simple | 2.10E-03 | -4.780 | -6.34 | mice2bgc | |
| [2] | models comparison | | sum of LL | p+ | log10(s+) | constrained LL | likelihood ratio | p-value |
| Constrained (H0): same s+, p+ | **human3bgc=fly3=mice3bgc** | | **-18.72** | **1.05E-03** | **-4.994** | **-431.44** | **825.43** | <1E-16 |
| | **human3bgc=fly3** | | **-12.38** | **1.05E-03** | **-4.995** | **-340.67** | **656.59** | <1E-16 |
| | human3bgc=fly2 | | -12.38 | 8.25E-04 | -4.970 | -342.95 | 661.15 | <1E-16 |
| | human2bgc=fly3 | | -12.38 | 1.05E-03 | -4.995 | -84.76 | 144.76 | <1E-16 |
| | human2bgc=fly2 | | -12.38 | 8.25E-04 | -4.970 | -85.51 | 146.26 | <1E-16 |
| | **human3bgc=mice3bgc** | | **-11.76** | **2.70E-03** | **-5.000** | **-323.63** | **623.74** | <1E-16 |
| | human3bgc=mice2bgc | | -11.76 | 3.53E-03 | -5.000 | -313.98 | 604.44 | <1E-16 |
| | human2bgc=mice3bgc | | -11.76 | 2.55E-03 | -5.000 | -79.16 | 134.79 | <1E-16 |
| | human2bgc=mice2bgc | | -11.76 | 3.53E-03 | -5.000 | -76.06 | 128.60 | <1E-16 |
| | **fly3=mice3bgc** | | **-13.31** | **7.50E-05** | **-3.842** | **-94.75** | **162.89** | <1E-16 |
| | fly3=mice2bgc | | -13.31 | 7.50E-05 | -3.840 | -524.63 | 1022.64 | <1E-16 |
| | fly2=mice3bgc | | -13.31 | 7.50E-05 | -3.926 | -118.44 | 210.26 | <1E-16 |
| | fly2=mice2bgc | | -13.31 | 7.50E-05 | -3.923 | -613.76 | 1200.90 | <1E-16 |

D. Test whether $\gamma^+$ and $p^+$ differ across taxa using only SSWW changes for human and mouse

| [1] model | species | outgroup | demorgraphic model | $p^+$ | $\gamma^+$ | log-likelihood | abbr. | |
|---|---|---|---|---|---|---|---|---|
| Full model (H1) | **Human** | **chimpanzee** | **Complex** | **0.01** | **6.18** | **-5.42** | **human3bgc** | |
| | Human | chimpanzee | Simple | 0.01 | 4.80 | -5.42 | human2bgc | |
| | ***D.melanogaster*** | ***D.simulans*** | **Complex** | **0.02** | **4.92** | **-6.96** | **fly3** | |
| | *D.melanogaster* | *D.simulans* | Simple | 0.04 | 2.79 | -6.96 | fly2 | |
| | **Mouse** | **Rat** | **Complex** | **0.01** | **1.29** | **-6.34** | **mice3bgc** | |
| | Mouse | Rat | Simple | 0.02 | 1.11 | -6.34 | mice2bgc | |
| [2] | models comparison | | sum of LL | $p^+$ | $\gamma^+$ | constrained LL | likelihood ratio | p-value |
| Constrained (H0): same gamma, p+ | **human3bgc=fly3=mice3bgc** | | **-18.72** | **0.05** | **1.26** | **-2266.67** | **4495.88** | **<1E-16** |
| | **human3bgc=fly3** | | **-12.38** | **0.06** | **1.00** | **-52.69** | **80.62** | **<1E-16** |
| | human3bgc=fly2 | | -12.38 | 0.06 | 1.00 | -90.94 | 157.12 | <1E-16 |
| | human2bgc=fly3 | | -12.38 | 0.06 | 1.00 | -45.22 | 65.67 | 5.5E-15 |
| | human2bgc=fly2 | | -12.38 | 0.06 | 1.00 | -65.69 | 106.62 | <1E-16 |
| | **human3bgc=mice3bgc** | | **-11.76** | **0.01** | **3.31** | **-124.80** | **226.08** | **<1E-16** |
| | human3bgc=mice2bgc | | -11.76 | 0.02 | 1.66 | -72.08 | 120.63 | <1E-16 |
| | human2bgc=mice3bgc | | -11.76 | 0.00 | 301.30 | -34.60 | 45.68 | 1.2E-10 |
| | human2bgc=mice2bgc | | -11.76 | 0.02 | 1.00 | -20.80 | 18.07 | 1.2E-04 |
| | **fly3=mice3bgc** | | **-13.31** | **0.05** | **1.31** | **-2231.12** | **4435.63** | <1E-16 |
| | fly3=mice2bbc | | -13.31 | 0.05 | 1.00 | -1992.65 | 3958.68 | <1E-16 |
| | fly2=mice3bgc | | -13.31 | 0.06 | 1.00 | -2784.95 | 5543.28 | <1E-16 |
| | fly2=mice2bgc | | -13.31 | 0.06 | 1.00 | -2559.72 | 5092.83 | <1E-16 |

[1]The top panel denotes the unconstrained model where each species is allowed to have its own $s^+$ (or $\gamma^+$) and $p^+$. The $s^+$ (or $\gamma^+$) and $p^+$ columns denote the maximum likelihood estimates (MLEs) of these parameters. The "abbr." gives the abbreviation for this demographic model used in the lower portion of the table. For example, "human3mac" means human Complex model using macaque as outgroup species.

[2]The bottom panel denotes the constrained model where $s^+$ (or $\gamma^+$) and $p^+$ were constrained to be the same across taxa. For example, "human3=fly3=mice3" means we constrained Complex models of human, *D. melanogaster* and mouse to have the same $s^+$ (or $\gamma^+$) and $p^+$. "Sum of LL" denotes the sum of the log-likelihoods across species for the full models listed in the second column. "Constrained LL" denotes the log-likelihood of the constrained model listed across the relevant species. "Likelihood ratio" denotes the difference in log-likelihood between the full and constrained models. *P*-values assume that twice the likelihood ratio is asymptotically distributed following a chi-square distribution.
